# Transposable elements may enhance antiviral resistance in HIV-1 elite controllers

**DOI:** 10.1101/2023.12.11.571123

**Authors:** Manvendra Singh, Sabrina M. Leddy, Luis Pedro Iñiguez, Matthew L. Bendall, Douglas F. Nixon, Cédric Feschotte

## Abstract

Less than 0.5% of people living with HIV-1 are elite controllers (ECs) - individuals who have a replication-competent viral reservoir in their CD4^+^ T cells but maintain undetectable plasma viremia without the help of antiretroviral therapy. While the EC CD4^+^ T cell transcriptome has been investigated for gene expression signatures associated with disease progression (or, in this case, a lack thereof), the expression and regulatory activity of transposable elements (TEs) in ECs has not been explored. Yet previous studies have established that TEs can directly impact the immune response to pathogens, including HIV-1. Thus, we hypothesize that the regulatory activities of TEs could contribute to the natural resistance of ECs against HIV-1. We perform a TE-centric analysis of previously published multi-omics data derived from EC individuals and other populations. We find that the CD4^+^ T cell transcriptome and retrotranscriptome of ECs are distinct from healthy controls, treated patients, and viremic progressors. However, there is a substantial level of transcriptomic heterogeneity among ECs. We categorize individuals with distinct chromatin accessibility and expression profiles into four clusters within the EC group, each possessing unique repertoires of TEs and antiviral factors. Notably, several TE families with known immuno-regulatory activity are differentially expressed among ECs. Their transcript levels in ECs positively correlate with their chromatin accessibility and negatively correlate with the expression of their KRAB zinc-finger (KZNF) repressors. This coordinated variation is seen at the level of individual TE loci likely acting or, in some cases, known to act as *cis*-regulatory elements for nearby genes involved in the immune response and HIV-1 restriction. Based on these results, we propose that the EC phenotype is driven in part by the reduced availability of specific KZNF proteins to repress TE-derived *cis*-regulatory elements for antiviral genes, thereby heightening their basal level of resistance to HIV-1 infection. Our study reveals considerable heterogeneity in the CD4^+^ T cell transcriptome of ECs, including variable expression of TEs and their KZNF controllers, that must be taken into consideration to decipher the mechanisms enabling HIV-1 control.

## Introduction

HIV-1 infection remains a major viral pandemic with an estimated 39 million people currently living with HIV-1 (PLWH), the majority of whom live in Sub-Saharan Africa**^1,2^**. Despite the availability of antiretroviral drug therapy (ART), there were more than 1.3 million new infections in 2022 alone**^3^**. There have been intense efforts to develop an effective vaccine and cure to HIV-1; in fact, over the past decade, several PLWH appear to have been cured**^4^**. Some received transplants with donor *CCR5Δ32* material rendering their cells resistant to R5 HIV-1 infection**^5–7^**. There are also post-ART controllers in whom no proviral integrations were detected for several years, suggesting a natural ability for viral maintenance**^8,9^**. Early in the HIV-1 pandemic, longitudinal studies showed that untreated HIV-1 infection led to a progressive loss of CD4^+^ T cells, but the rate of CD4^+^ T cell decline varied from person to person**^10^**. A small number of PLWH had no clinical symptoms and no CD4^+^ T cell decline. These long-term non-progressors were dubbed “elite controllers’’ (EC). Some EC developed low level viremia and CD4^+^ T cell loss over time, while others maintained viremic control long-term and became known as “exceptional elite controllers”**^11–13^**. These exceptional ECs include those who appear to have cleared the virus completely**^8,9^**. The control of viral replication in PLWH varies widely between individuals. However, our understanding of the molecular mechanisms underlying this range of antiviral resistance remains limited.

Seminal genome-wide association studies of the EC phenotype identified cell-mediated immunity**^14^** and the HLA-B region as major determinants of viral control**^15,16^**. Subsequent studies of NK cell receptors identified a number of innate immune responses as additional contributors to low level viremia**^17^**. More recent studies have characterized proviral reservoirs in ECs**^9,18,19^**. Compared to PLWH-on-ART, intact proviral sequences from ECs were found to have integrated at distinct sites in the human genome, preferentially located in centromeric satellite DNA or in KRAB-containing zinc finger (KZNF) genes on chromosome 19**^9^**, which are associated with heterochromatin features**^20^**. Moreover, the integration sites of intact proviral sequences from ECs appear further from transcriptional start sites and accessible chromatin of the host genome than in PLWH-on-ART and were enriched in repressive chromatin marks**^9,19^**. In contrast, defective proviruses in ECs were commonly located in permissive genic euchromatin positions**^9,18^**.

TEs are mobile DNA elements that can replicate and insert themselves into different locations within the host genome. Nearly half of the human genome is composed of these elements, though most TE loci have long since lost the ability to self-propagate due to mutational decay and/or epigenetic silencing from repressive factors like KZNFs**^21–23^**. Despite their immobilization, many human TEs still show transcriptional activity and regulatory capabilities, the consequences of which are the subject of intense investigation. Multiple studies have documented cases of co-option of TE-derived *cis*-regulatory elements for host gene regulation, including the control of innate immunity genes**^21,24–28^**. For example, we have previously demonstrated that multiple elements from the human endogenous retroviruses (HERV) family MER41 function as interferon-inducible enhancers triggering the activation of innate immunity genes upon infection, including those involved in restricting HIV-1**^29^**. Furthermore, elements from the LTR12C and LTR7 HERV families are known to serve as promoters or enhancers for human genes encoding restriction factors against HIV-1 replication, specifically GBP2/5 and APOBEC3G/H, respectively**^30–33^**. Thus, interindividual variation in the *cis*-regulatory activity and expression profiles of these and other TEs holds the potential to directly influence viral susceptibility, including susceptibility to HIV-1 infection**^34–36^**. Here, we begin testing the hypothesis that the EC phenotype may be driven, in part, by the increase in TE *cis*-regulatory activity and their resulting capacity to influence immune gene expression.

To examine this, we investigate whether ECs can be distinguished by differential regulation of TEs from ART-treated people living with HIV (PLWH-on-ART), treatment-naive viremic progressors (VPs), HIV-1 uninfected healthy controls (HCs), as well as from each other (EC vs. EC). Through parallel transcriptomic and epigenomic analyses on publicly available CD4^+^ T cell data**^9,37–39^**, we find that the TE transcriptome (also known as the retrotranscriptome) of ECs is distinct from HCs and VPs. Furthermore, we find that ECs are transcriptionally heterogenous and can be divided into four clusters distinguished by their expression of innate immune genes and TE families. In a subset of ECs, we identify an increase in the chromatin accessibility of specific TE loci which correlates with higher expression of nearby HIV-1 restriction factors and other immune-related genes in ECs compared to HCs. Finally, we observe that transcript levels of KZNFs in EC CD4^+^ T cells negatively correlate with that of the TE families they are predicted to transcriptionally repress, suggesting that interindividual genetic and/or epigenetic variation of KZNFs may underlie differential TE regulation in ECs. We speculate that derepression of certain TEs could boost their *cis*-regulatory activity on antiviral genes thereby enhancing the immune response to HIV-1 infection. Overall, our findings support the notion that the genomic activation of specific TE loci may contribute to HIV-1 resistance.

## Results

### The EC (retro)transcriptome is distinct from healthy controls, PLWH-on-ART, and viremic progressors

In order to identify unique transcriptomic and retrotranscriptomic (defined as TE transcripts) features of ECs, we first analyzed RNA-Seq data from a study investigating the role of the HIV-1 coreceptor CCR5 in the EC phenotype, which was generated from activated CD4^+^ T cells of ECs (n=4) and HCs (n=5)**^39^**. The differential expression of immune-related genes between ECs and HCs has been documented**^38,40^**, but the EC retrotranscriptome has yet to be profiled. We analyzed this dataset to identify the genes and TE families that were differentially expressed between the two populations (Table S1). The differential expression of genes and TEs was similar in magnitude, with maximum log2 fold changes of significance around ±10 (Figure 1A).

**Figure 1.**
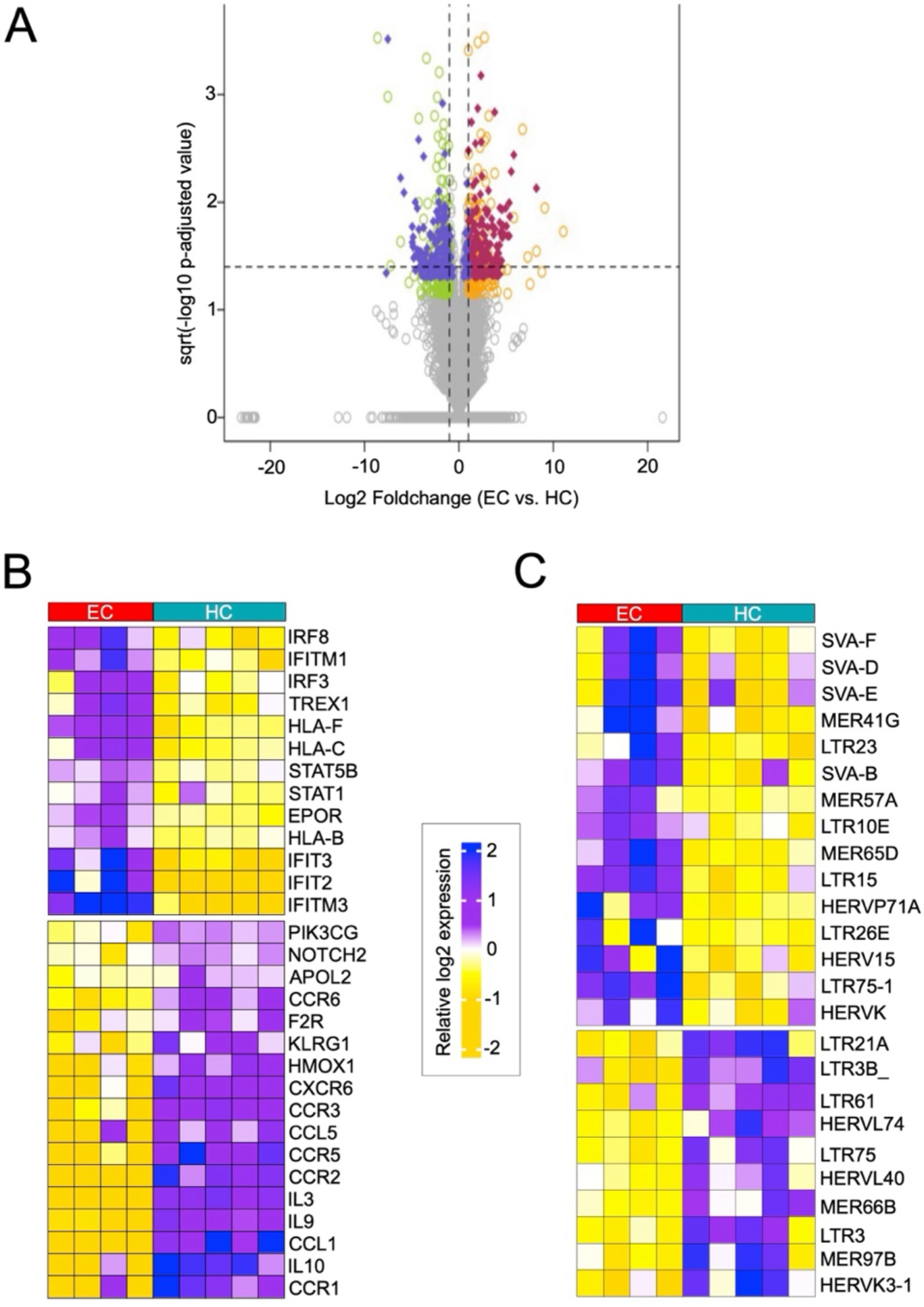
Differential (retro)transcriptomic profiles in ECs vs. HCs. **A.** Volcano plot illustrating the differentially expressed TEs and genes between ECs (n=4) and HCs (n=5). Coloration is based on increased or decreased expression of genes (orange and green, respectively) and TEs (red and purple, respectively). Total detected genes and TE loci are plotted by log2-transformed fold change. Statistical significance given in the form of sqrt −log10 adjusted p-value, calculated by Wilcoxon rank sum test with Bonferroni correction. Data source: <u>Gonzalo-Gil et al., 2019</u>. **B.** Heatmap displaying the expression of the top differentially expressed genes in CD4^+^ T cells of ECs (n=4; red bar) vs. HCs (n=5; blue bar). Heatmap coloration is based on the z-score distribution from −2 to 2 (gold to purple; low to high expression). Data source: <u>Gonzalo-Gil et al., 2019</u>. **C.** Heatmap displaying the expression of the top differentially expressed TE families in CD4^+^ T cells of EC (n=4; red bar) vs. HCs (n=5; blue bar). Heatmap coloration is based on the z-score distribution from −2 to 2 (gold to purple; low to high expression). Data source: <u>Gonzalo-Gil et al., 2019</u>.

Among the differentially expressed genes, we noted the lower expression of factors known to facilitate HIV-1 entry in ECs, including CCR2/3/5/6 and CXCR4 receptors (Figure 1B), consistent with the study from which the RNA-seq data originate**^39^**. Furthermore, multiple HLA genes were more highly expressed in ECs, including HLA-B, −C, and −F (Figure 1B). We and others have shown differences in antiviral restriction factor expression between PLWH and HCs based on their HLA genotypes**^15,41,42^**, and certain HLA-B alleles have been strongly associated with elite control in previous GWAS studies**^16,17^**. We also observed elevated transcript levels for several restriction factor-encoding genes (IFIT and IFITMs) and pro-inflammatory factors (STAT1, IRF8) known to be activated in response to viral infections, including HIV-1**^43–45^**, which may contribute to the EC phenotype. For example, the immune transcription factor IRF8 is highly expressed in ECs relative to HCs (Figure 1B). Low levels of IRF8 have been previously associated with adverse neurological outcomes in PLWH-on-ART**^46^**. This higher expression may be yet another protective aspect of elite control. Thus, the distinct immune gene expression profile we observe in ECs is largely consistent with previous studies and brings further support to the idea that enhanced innate immunity contributes to HIV-1 restriction in ECs**^47–50^**.

The most differentially expressed TE subfamilies were dominated by primate-specific TEs – specifically SVA and LTR/ERV elements – known to harbor complex *cis*- regulatory sequences (Figure 1C). For example, numerous previous studies have found that SVA and HERVK elements are frequently co-opted as *cis*-regulatory elements in human gene regulatory networks, including those involved in embryonic genome activation**^51,52^** and cell type identity**^53–56^**. Similarly, many LTR10 elements carry binding sites for p53**^57^** and enhancers activated in cancer**^58,59^**. Finally, MER41, LTR26, and MER57 – from which certain subfamilies have higher expression in ECs compared to HCs (Figure 1C) – are known to be enriched for STAT1 binding sites and frequently behave as interferon-inducible enhancers associated with innate immunity genes**^29^**.

Conversely, the TE families with lower expression in ECs are also enriched for LTR/ERV subfamilies (Figure 1C), suggesting broad differences in the regulatory landscape of CD4^+^ T cells in ECs compared to HCs.

Knowing that ECs are distinct from HCs, we aimed to validate that these differences were not solely attributed to the presence of HIV-1 in one group but not the other. Thus, we next juxtaposed ECs with PLWH-on-ART. We compared TE family expression in CD4^+^ T cell subsets of ECs (n=12) and PLWH-on-ART (n=3) using RNA-seq data from a previous study that investigated EC-specific proviral integration patterns (Table S2)**^9^**. Consistent with our EC vs. HC comparisons, MER41 and SVA subfamilies were more highly expressed in ECs than in PLWH-on-ART (Figure S1), in addition to ERV subfamilies LTR12, LTR13, and THE1B.

To ensure that these findings were not confounded by the presence of antiretroviral drug treatment, we performed a more extensive comparison between the (retro)transcriptomes of untreated ECs (n=19) and treatment-naïve viremic progressors (VPs; n=8). This data came from a study identifying differentially expressed transcripts in the donors’ peripheral blood mononuclear cells (PBMCs)**^38^**. After identifying the most variable genes and TEs (Table S3), a principal component analysis (PCA) was conducted to evaluate the level of transcriptional separation between the ECs and VPs (Figure 2A). Approximately 1900 genes and 100 TE families segregated the EC and VP samples on the first three principal components, indicating that ECs have distinct transcriptomic and retrotranscriptomic profiles from VPs (Figure 2B). In fact, when conducting a separate PCA that included the HC samples from the same study, the ECs appear to be even more distinct from the VPs than from the HCs (Figure S2A&B).

**Figure 2.**
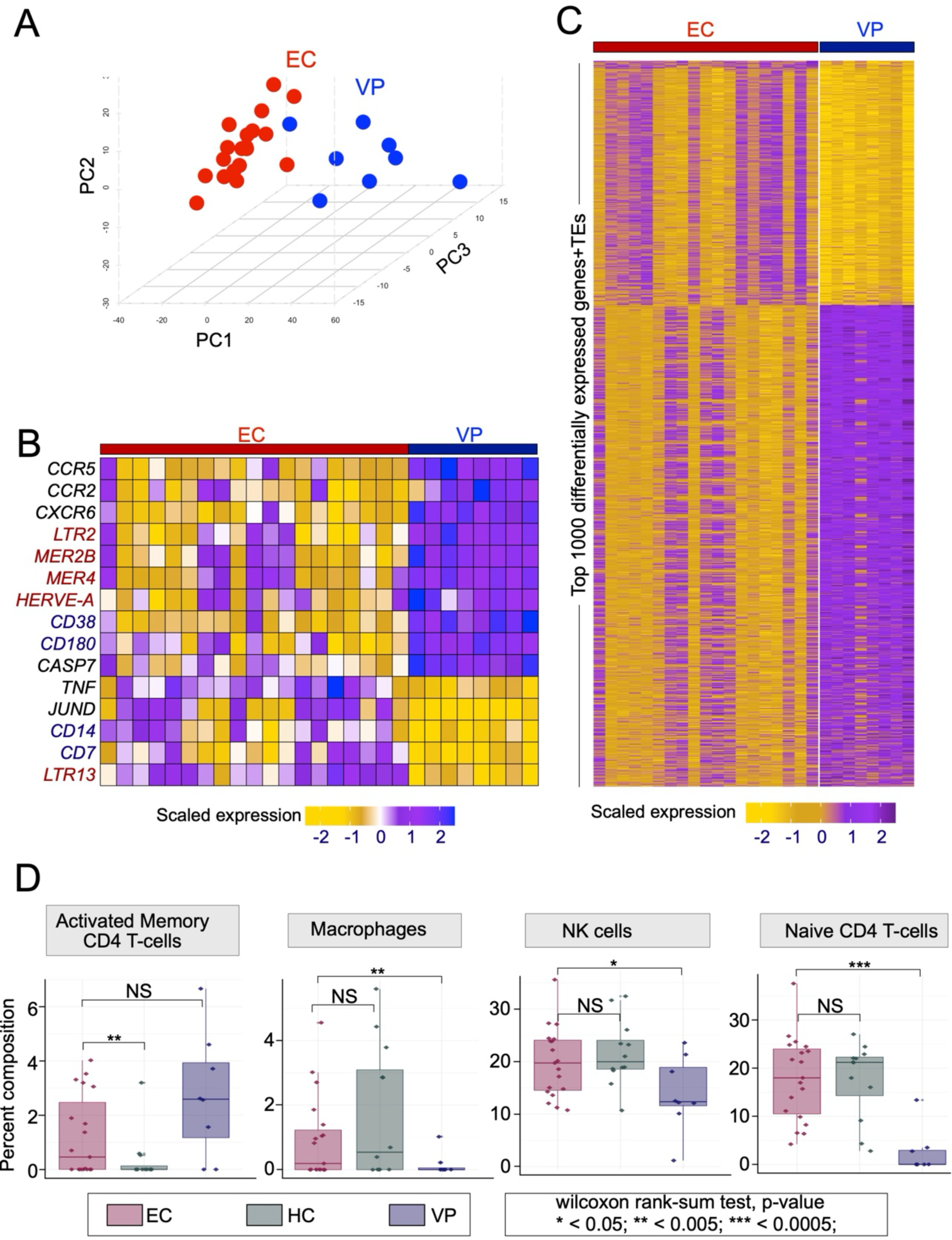
Differential (retro)transcriptomic & immune cell profiles in ECs vs. VPs. **A.** PCA triplot from PBMCs of ECs (red) and VPs (blue), based on the most variably expressed genes and TE families. Data source: <u>Zhang et al., 2018.</u> **B.** Heatmap of z-scaled expression (log2 TPM) from select gene/TE sets between ECs and VPs. On the y-axis, immune genes are in black, leukocyte surface markers are in blue, and TEs are in red. Data source: <u>Zhang et al., 2018.</u> **C.** Heatmap displaying the z-scaled expression (log2 TPM) of genes and TEs distinguishing EC and VP RNA-seq samples. Every row denotes a gene or TE element. Data source: <u>Zhang et al., 2018.</u> **D.** Box plots for leukocyte population of interest, identified via deconvolution analysis of PBMC RNA-seq data. Statistical significance determined by Wilcoxon rank-sum tests. Data source: <u>Zhang et al., 2018</u>.

Immune signatures in our EC vs. VP comparison were generally consistent with previous comparisons of ECs to other groups, including the notably lower expression of CCR5 in ECs (Figure 2C), even though no EC donor harbored the *CCR5Δ32* deletion**^38,39^**. Additional notable differences between the ECs and VPs included high expression of TNF in ECs, a cytokine with a crucial role in the immune response (Figure 2C). Furthermore, we observed differential expression of numerous leukocyte surface markers: CD14 (macrophage marker) and CD7 (effector CD8+ T cell marker) had higher expression in ECs, while expression of activated immune cell markers CD38 and CD180 was higher in VPs. For the retrotranscriptome, we noted the higher expression of the LTR13 subfamily in ECs compared to VPs, consistent with our EC vs. PLWH-on-ART analysis and of note because LTR13 is a subfamily known to be enriched for STAT1 binding**^29^**. Taken together, these analyses suggest that the ECs have a (retro)transcriptomic profile distinct from HCs, VPs, and PLWH-on-ART characterized by elevated levels of immune-responsive genes and specific TEs.

Having focused thus far on identifying transcriptional differences, we next aimed to determine if ECs, VPs, and HCs had distinct immune cell compositions as well. With the abovementioned PBMC RNA-seq dataset**^38^**, we conducted deconvolution analyses through which we were able to distinguish transcriptional signatures of 19 immune cell types. In line with the above analyses, ECs, VPs, and HCs had distinct immune cell compositions (Figure S3). Most significantly, ECs had a higher proportion of macrophages, naïve CD4^+^ T cells, and NK cells compared to VPs, and a higher proportion of activated memory CD4^+^ cells compared to HCs (Figure 2D). Thus, the transcriptomic differences observed between ECs and other populations may be partially driven by variation in immune cell type composition.

### CD4^+^ T cells from ECs can be split into four functionally distinct clusters

In our initial differential expression analyses, we observed consistent expression patterns among HCs and VPs, but found notable heterogeneity between ECs (Figure 1B, 1C, 2B, 2C, & S2B). To further explore this phenomenon and identify markers that capture the transcriptomic and retrotranscriptomic heterogeneity of CD4^+^ T cells in ECs, we analyzed a larger group of ECs using two previously published RNA-Seq datasets of CD4^+^ T cell subtypes**^9,37^**including naïve, central memory (CM), effector memory (EM), transitional memory (TM), and total CD4^+^ T cells. The first study used RNA-Seq to characterize the proviral reservoir of 12 ECs compared to 3 PLWH-on-ART**^9^**. The second study investigated mechanisms for viral persistence in 15 EC individuals**^37^**. We integrated the RNA-seq data from both studies (a cumulative 128 EC samples; Table S4) to increase our power to potentially classify ECs into subgroups. We then performed unbiased clustering on the scaled & normalized RNA-seq data by feeding the first five principal components to the graph-based k-nearest neighbors (KNN) algorithm (see Methods). This analysis revealed four distinct clusters (Figure 3A). Importantly, the samples did not cluster by patient cohort or study of origin, thereby ruling out batch effects as the drivers of the clustering (Figure S4A&B). Furthermore, the addition of the sequenced ART-treated samples (n=15) from one of the two studies of origin did not disrupt the number or separation of the EC clusters**^9^**. To examine whether the clustering could instead be explained by patient ancestry, we next visualized the samples by ancestry as inferred by variant comparisons with HapMap (Figure S4C)**^60^**. All clusters were heterogeneous for patient ancestry. Finally, we visualized the samples by cellular subtype, as identified in the original studies, to assess whether the clustering can be explained by CD4^+^ T cell subtype composition (Figure S4D). Clusters 1 and 2 were essentially indistinguishable by cell type composition, whereas Clusters 3 and 4 showed an overrepresentation of TM/EM and naïve/CM cell types, respectively (Figure 3B). Thus, cell subtype composition could only partially explain the clustering.

**Figure 3.**
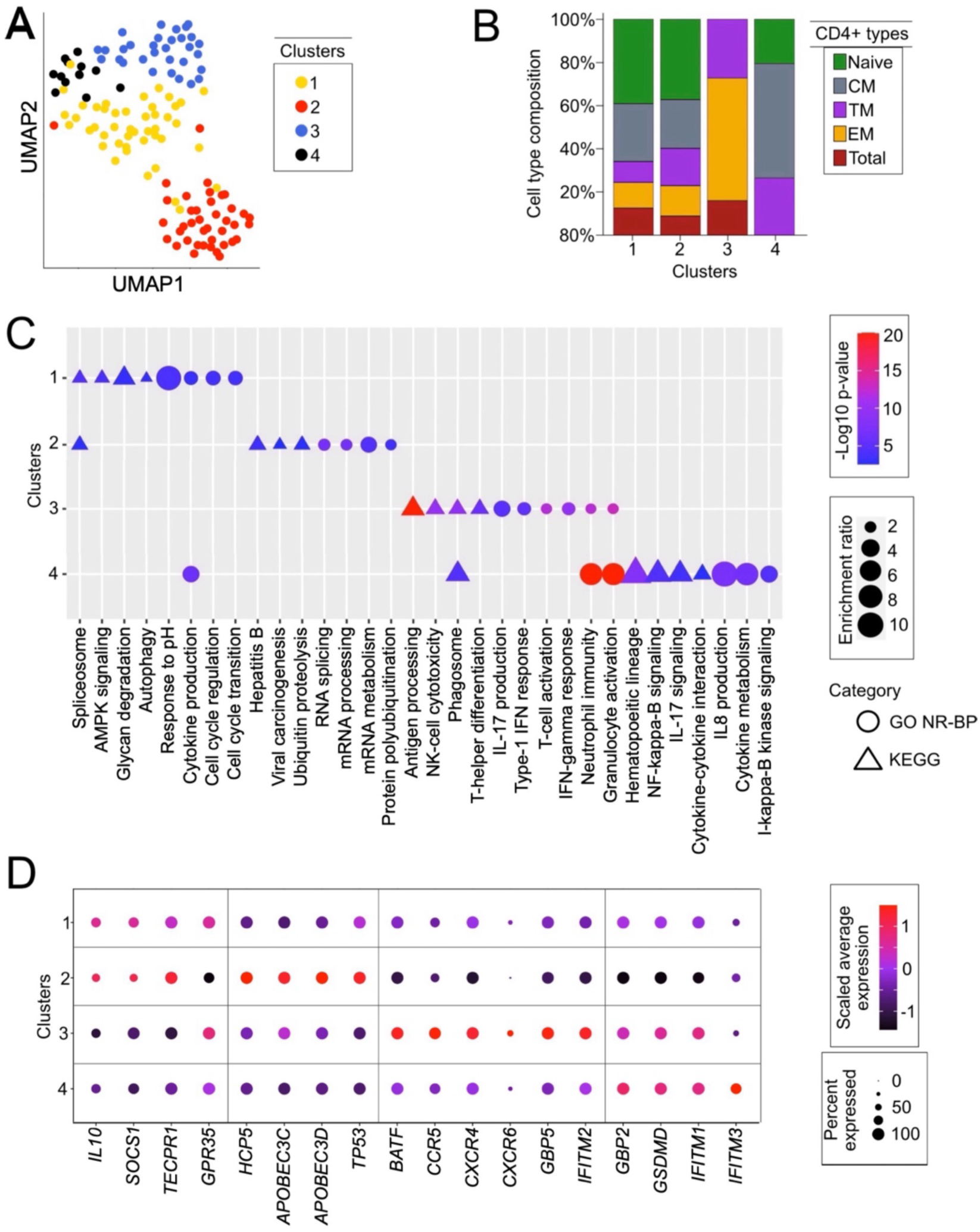
EC CD4^+^ T cells can be grouped into four distinct clusters. **A.**UMAP plot of the four clusters of EC CD4^+^ T cell subtypes (N=128) using KNN graph construction on bulk RNA-seq data, based on the Euclidean distance in PCA space (see Methods). Every point is a CD4^+^ T cell RNA-seq sample, colored by cluster assignment. Data sources: <u>Jiang et al., 2020</u> and <u>Boritz et al.,</u> <u>2016</u>. **B.** Stacked barplot displaying the composition of CD4^+^ T cell subtypes (naïve, CM, TM, EM, total) in each of the four clusters. Data sources: <u>Jiang et al., 2020</u> and <u>Boritz et al., 2016</u>. **C.** Gene ontology biological process (GO NR-BP; ●) and KEGG pathway (KEGG; ▴) delineation of the four EC clusters using WebGestalt**^70^**. Data derived from differential expression analysis of the EC clusters, using the significant DEGs (p- value < 0.05) as each cluster’s respective gene list. For each of the four EC clusters, the highest ranked GO terms and KEGG pathways by adjusted p-value are shown. ‘Enrichment ratio’ refers to the number of observed genes divided by the number of expected genes from each GO or KEGG category in the cluster’s gene list. Data sources: <u>Jiang et al., 2020</u> and <u>Boritz et al., 2016</u>. **D.** Dot plot illustrating the scaled expression of selected genes related to HIV-1 replication in the four EC clusters. Coloration represents the log2-transformed expression scaled to their transcriptome and averaged across the cluster’s samples, from lower (blue) to higher (red) expression. The size of the dots is directly proportional to the percent of samples expressing the given gene in a given cluster. Data sources: <u>Jiang et al., 2020</u> and <u>Boritz et al., 2016</u>.

To functionally characterize these four clusters, we performed pathway and gene ontology analyses on each cluster using gene lists that defined the individual clusters, extracted by differential expression analysis of the cluster’s samples versus the samples of the remaining three clusters (Table S5). Using both the KEGG pathway database**^61^** and the biological process aspect from the Gene Ontology Consortium**^62^**, we were able to identify distinct pathways and ontological enrichments for each of the four clusters (Figure 3C & Table S6). Genes related to cell turnover, autophagy, and cell cycle regulation were overrepresented in Cluster 1. Genes related to RNA processing and splicing were overrepresented in Cluster 2. Clusters 3 and 4 had some overlap, as both contained markers of immune function, but with some distinguishing features. Cluster 3 showed enrichment for genes regulating T cell activation and proliferation, whereas Cluster 4 was enriched for genes involved in neutrophil–T cell interactions and cytokine production. The enrichment of TM and EM CD4^+^ T cells in Cluster 3 aligns with the overrepresentation of pathways related to T cell activation and proliferation. Similarly, the enrichment of naïve and CM CD4^+^ T cells in Cluster 4 aligns with the overrepresentation of pathways related to neutrophil-T cell interactions and cytokine production, reflecting their involvement in pathogen surveillance and immune cell communication. Thus, we concluded that EC transcriptomes can be classified into four distinct clusters defined by the enrichment of specific biological functions.

With a better understanding of the functional delineation between the four EC clusters, we next wanted to examine whether cluster-specific differences could reveal different mechanisms of HIV-1 resistance. To accomplish this, we focused on differentially expressed genes that encode factors known to directly modulate HIV-1 infection. Cluster 2, which is the most distinct in our UMAP (Figure 3A), had the most distinct gene expression profile, characterized by the relatively low expression of HIV-1 coreceptors CCR5 and CXCR4 and the high expression of innate immunity genes like HCP5 and APOBECs (Figure 3D)**^32,63,64^**. Additionally, Cluster 2 showed high expression of TP53, known to be involved in heterochromatin expansion and inhibition of early HIV- 1 replication**^65,66^**. Within the other clusters, we found that Clusters 3 and 4 each expressed at least one restriction factor (IFITM1, IFITM2, IFITM3, GBP2, GBP5, etc.) at a significantly higher level than Clusters 1 and 2 (Figure 3D).

Taken together, these data indicate that HIV-1 resistance in ECs does not stem from a single factor but reflects a complex phenotype driven by an interplay between the expression of HIV-1-restrictive and HIV-1-facilitative factors, in contrast to previous observations suggesting that the majority of the EC phenotype was driven by HLA-B or KIR-mediated cellular immune effects**^67–69^**. Furthermore, our clustering analysis suggests that it is possible to stratify this phenotype according to gene expression signatures, including but not limited to the expression of key factors regulating HIV-1 infection and replication.

### Restriction factor expression in ECs may be driven by more accessible retroelements

Previous studies have implicated TEs as direct regulators of interferon-responsive gene expression upon viral infection, including antiviral factors**^29,31^**. Having determined that both TEs and HIV-1 restriction factors are differentially expressed in ECs compared to HCs, VPs, and PLWH-on-ART, we next aimed to determine whether changes in TE expression and *cis*-regulatory activity could be correlated to changes in innate immune gene expression. We used paired ATAC-seq – which measures chromatin accessibility – and RNA-seq datasets for ECs (n=4) and HCs (n=4)**^39^** to examine whether the chromatin accessibility of TE integrants located near genes correlated with the gene’s expression level in ECs, using a 10 kb range from the transcription start site as our cutoff. We identified over 250 TE-gene pairs where we observed such correlations, with increased TE accessibility correlating to increased gene expression in ECs compared to HCs (Table S7). Of these, approximately one third are annotated in relation to immune response and, interestingly, many of these involve TE families previously implicated in innate immunity and known to be bound by transcriptional activators of the innate immune response (Figure 4). For example, GPR35**^71^**, GSDMD**^72^**, and TECPR1**^73^** are three antiviral genes more highly expressed in ECs compared to HCs which are flanked by MER41 elements bound by STAT1 and STAT5 and marked by more accessible chromatin in ECs (Figure 4). Similarly, GBP2 and GBP5 are two guanylate binding proteins known to restrict HIV-1 entry and previously shown to be transcribed from LTR12C elements upon HIV-1 infection of primary CD4^+^ T cells**^31^**. We found that transcript levels for both GBP2 and GBP5 are moderately (but still significantly) elevated in ECs compared to HCs and observed that their respective LTR12C-derived promoters are marked by more accessible chromatin in ECs (Figure 4). The final case highlighted in Figure 4 is HCP5, a lncRNA which has been implicated in the innate immune response to several pathogens, including HIV-1**^74,75^**. We found that HCP5 expression is higher in ECs compared to HCs, correlated to the accessibility of its ERV3-derived promoter. Together these examples suggest that the upregulation of HIV-1 restriction factors and other antiviral genes in ECs may be driven by increased *cis*-regulatory activity of the nearby TEs controlling their expression.

**Figure 4:**
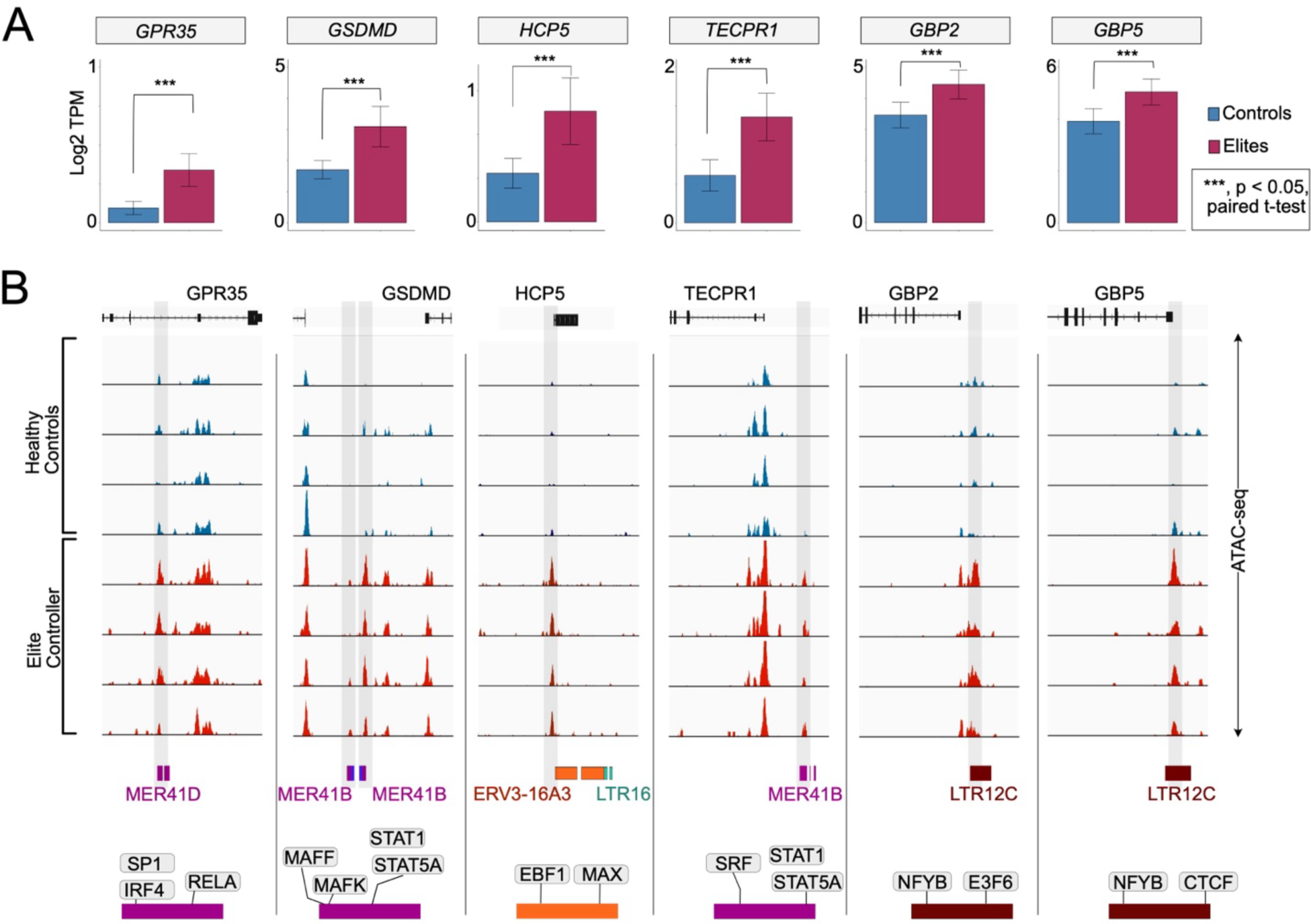
Induction of innate immune gene expression by proximal LTRs. **A.** Barplots showing the expression (log2 TPM) of selected differentially expressed innate immune genes in HCs (n=5) and ECs (n=4). P-value is calculated by paired student t-test. RNA- & ATAC-seq data source: <u>Gonzalo-Gil et al., 2019.</u> **B.** Integrative genome visualization (IGV) of normalized ATAC-Seq signal around the selected DEGs in Figure 5A between HC (n=4) and EC (n=4) CD4^+^ T cells. ATAC-seq peaks of interest are shaded in light gray. The proximal TE integrants are shown below the IGV graph, under which the encoded TF binding over the corresponding TE integrant(s) is also shown. RNA- & ATAC-seq data source: <u>Gonzalo-Gil et al., 2019.</u>

### EC clusters are characterized by increased expression of specific TE families, potentially mediated by KZNF derepression

Building upon the hypothesis that the activity of certain TE loci may enhance the expression of antiviral genes in ECs (Figure 4) and the knowledge that many of these antiviral factors are differentially expressed across the four EC clusters we identified (Figure 3), we hypothesized that the expression of certain TE families may vary across these clusters. To test this, we first compared TE transcript levels across the RNA-seq datasets used to define the four EC clusters**^9,37^**. For the samples in each cluster, we calculated TE family expression by averaging the transcript levels of all loci from a given family and comparing these average values between clusters. As we observed in our earlier comparisons of the EC clusters’ immune profiles (Figure 3D), we found that each EC cluster was characterized by a distinct TE expression profile. Figure 5A highlights a subset of differentially expressed TE families of particular interest. Cluster 1 was characterized by high expression level of the youngest LINE1 subfamilies in the human genome (L1HS, L1PA2, L1PA3)**^76^**. Cluster 2 was marked by high levels of LTR7/HERVH and MER41B, two subfamilies previously implicated in the regulation of antiviral factors**^29,33^**. Cluster 3 was marked by high LTR12C expression, consistent with the high level of GBP2 and GBP5 observed in some EC individuals (Figure 4) including those falling within Cluster 3 (Figure 3D). Finally, Cluster 4 was characterized by a high level of HERVL40 expression, a TE subfamily previously observed to be downregulated upon HIV-1 latency reversal**^77^**. Thus, each cluster displayed a unique profile of gene and TE expression (Figure S5).

**Figure 5.**
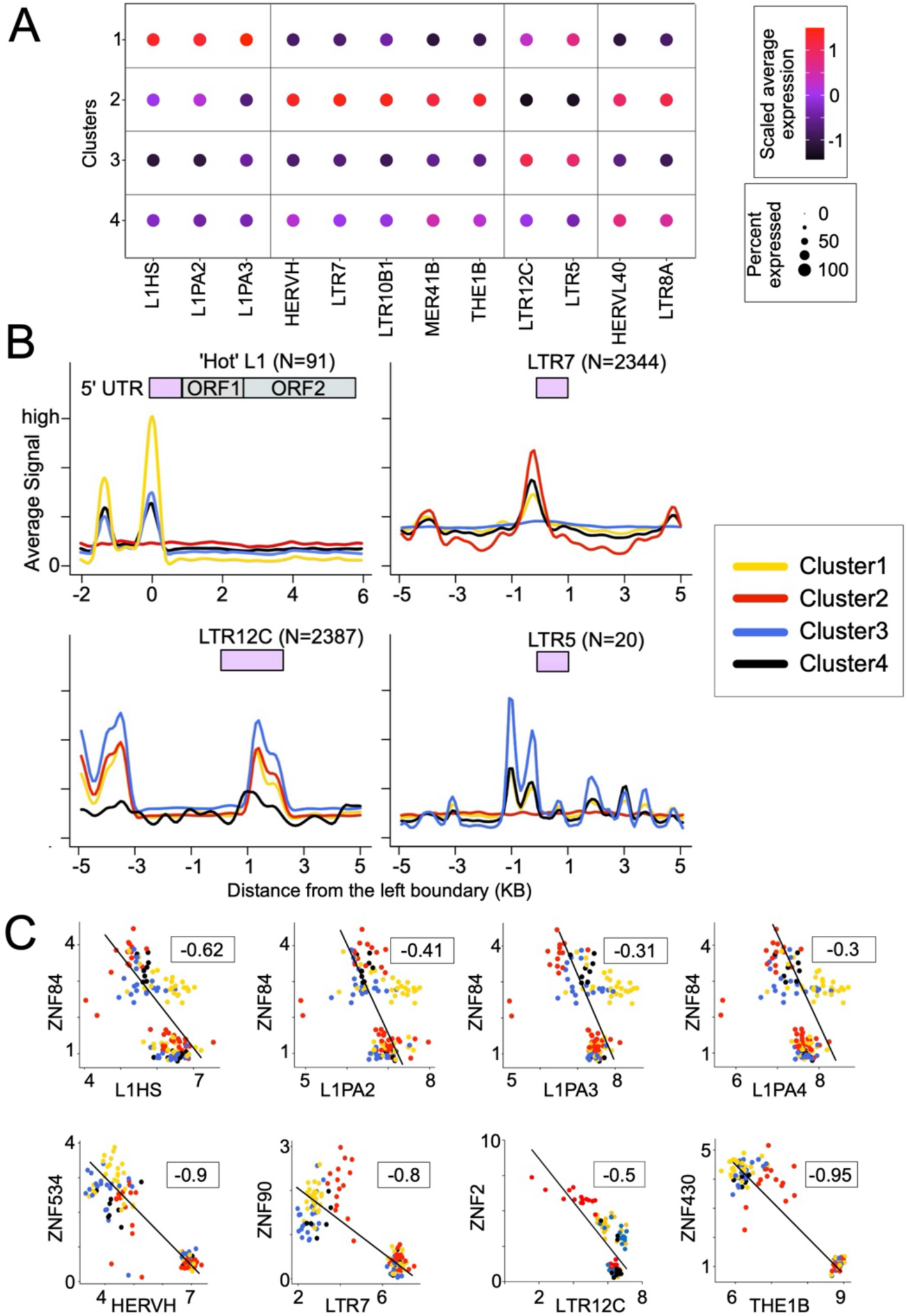
Characterization of TE expression in specific EC clusters and its correlation to KZNF repression. A. Dot plot illustrating the intensity and abundance of TE family expression (log2 CPM+1) across the four EC clusters. Coloration is scaled from lower (black) to higher (red) expression. Point size is directly proportional to the percent of samples expressing the TE family in the given cluster. Data sources: <u>Jiang et al., 2020</u> and <u>Boritz et al., 2016</u>. B. Line plots showing the distribution of averaged, normalized ATAC-seq signal over all loci in the selected TE families for the samples in each cluster. The ATAC-seq signal counts are calculated and normalized as mappable reads per million per 100 bp bins, in a +/- 5 kb genomic window at the elements’ left boundary in the human genome. For the L1 elements, only those that are intact and full-length are considered, denoted as ‘hot’ L1s for their recent transpositional activity. Lines are color coded by cluster. Data source: <u>Jiang et al., 2020</u>. C. Scatter plot showing scaled, log2-transformed expression of previously implicated TE families and KZNFs in the analyzed CD4^+^ T cells subtype EC RNA-seq data, cluster classified in Figure 5A. Linear regression analysis (black line) indicates the correlation between TE families and their targeting KZNF’s expression in EC samples. The rho value is obtained from pairwise-ranked correlation analysis. Data sources: <u>Jiang et al., 2020</u> and <u>Boritz et al., 2016.</u>

To examine whether the expression profiles of specific TE families in the EC clusters correlated with the chromatin accessibility of their canonical promoters, we analyzed CD4^+^ T cell ATAC-seq data produced in parallel with the RNA-seq data used in the EC clustering, available for a subset of the ECs (n=60)**^9^**. For the samples in each cluster, we calculated the averaged, normalized ATAC-seq signal over all loci of the TE family of interest and compared the average signal for each cluster. For most TE subfamilies, we found that the average chromatin accessibility profile across clusters correlated with their RNA expression profile (Figure 5B). For example, the promoter region of young L1 elements was more accessible in individuals from Cluster 1 where these elements are most highly expressed. Likewise, LTR7 elements were on average more accessible in individuals from Clusters 2 and 4 in which these elements are also most highly expressed (Figure 5A&B). Of those highlighted in Figure 5A, there were four TE subfamilies for which we could not correlate subfamily-wide average chromatin accessibility and cluster-specific expression: MER41B, LTR10B1, THE1B, and LTR8A. For these subfamilies, it appears that high expression in a certain cluster is driven by a small subset of particularly active loci, limiting visibility at the family level.

What could drive the differential chromatin accessibility and expression of these TEs across ECs? We hypothesized that KRAB-containing zinc finger proteins (KZNFs) may be involved since they are known to bind directly to specific TE subfamilies through their zinc finger domain, recruit the KAP1 corepressor through their KRAB domain, and in turn attract silencing factors such as histone deacetylases and methyltransferases to nucleate repressive chromatin at the bound TE loci**^78^**. To test this idea, we performed a pairwise correlation analysis of TE families and KZNFs from the transcriptomes of the EC samples used for the initial clustering analysis. With over 300 annotated KZNFs in the human genome, the repressive relationship between these proteins and TEs is thought to be highly specific**^79^**, and as such we predicted that KZNFs and their target TE subfamilies would show a negative correlation in expression across individuals.

Visualizing these correlation analyses by EC cluster, we observed distinct anti-correlative KZNF-TE expression patterns (Figure S6A), suggesting that each cluster is characterized by a unique combination of anticorrelated KNZF-TE pairs. Focusing on the TE families that stood out in our earlier analyses, we found multiple KZNFs whose expression was anticorrelated to that of the transcriptionally elevated TE families they are known to target (Figure 5C). For example, EC expression of ZNF84 – which is known to bind and repress young L1 subfamilies**^80^**– was significantly anticorrelated to the expression of the same L1 subfamilies. Similarly, ZNF534 was anticorrelated with HERVH/LTR7, ZNF430 with THE1B, and ZNF2 with LTR12C. Binding between all of the aforementioned TE subfamilies and their respective KZNFs was confirmed by ChIP-exo (Figure S6B)**^80,81^**. Notably, the majority of these KZNFs have significantly lower expression in ECs compared to treatment-naïve VPs (Figure S6C). Thus, the expression of these KZNFs may explain the variable expression of these TEs across EC individuals and between HIV-infected populations. Based on these analyses, we believe that the heterogeneous EC phenotype may be driven in part by the reduced ability of KZNFs to repress specific TEs that serve as *cis*-regulatory elements for nearby antiviral genes, thereby heightening their resistance to HIV-1 infection.

## Discussion

In this study, we have begun exploring the hypothesis that TEs, which are known to modulate the innate immune response and directly regulate restriction factors, could contribute to the elite control of HIV-1. Through bulk retro-transcriptomic analyses of available CD4^+^ T cell RNA-seq data, we determined that the TE expression profile of ECs is distinct from that of PLWH-on-ART, VPs, and HCs. There was also considerable transcriptomic heterogeneity among the set of EC individuals analyzed here, despite being a relatively small cohort (n=49, combining four different studies)**^9,37–39^**. Unsupervised clustering and principal component analysis of the ECs’ most variable genes and TEs revealed four clusters of ECs, each characterized by different gene ontologies and pathway enrichments, differential expression of known HIV-1 restriction factors, and unique TE expression profiles. Further analyses integrating parallel ATAC- seq data revealed that several innate immune genes with increased expression in ECs were flanked by *cis*-regulatory TEs marked with higher chromatin accessibility in ECs. This suggests that changes in the chromatin states of these elements contribute to the EC-specific upregulation of these factors, which in turn may contribute to their HIV-1 resistance phenotype. To begin understanding the mechanism driving chromatin changes at these TEs, we investigated the expression of KZNFs across the same EC cohort – genes which are known to encode proteins that repress TE expression and *cis*- regulatory activity**^22,82^**. We found extensive variation in KZNF expression levels across CD4^+^ T cells of EC individuals and striking anticorrelation with the TE families they are known to target, suggesting that interindividual variation in KZNF expression may underlie variation in TE chromatin accessibility. Taken together, these data converge to a model (Figure 6) in which low expression of some KZNFs leads to elevated basal expression of innate immune genes and restriction factors in EC individuals through an increase in the *cis*-regulatory activity of TEs serving as promoters or enhancers for these genes. This model introduces a new possible mechanism underlying the elite control phenotype.

**Figure 6.**
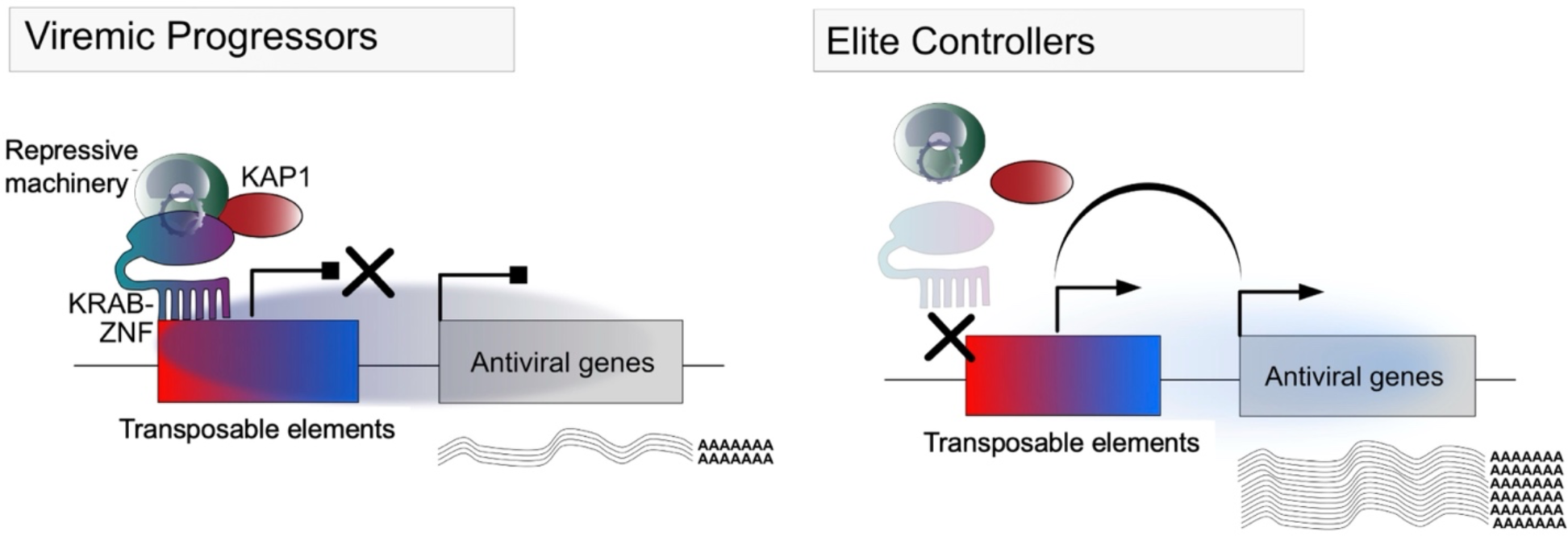
The interplay of KRAB-ZNFs and TEs regulates the expression of proximal antiviral genes.

Each step in the model will require experimental work to be validated. First and foremost, it will be important to confirm that the TEs identified as exhibiting increased transcript levels and accessibility in ECs are indeed boosting the innate immune response and control of HIV-1 in these individuals. We note that these TEs include HERV families that have been previously implicated in controlling the human innate immune response. For example, we found that the MER41 family has increased expression in a subset of ECs (Figure 5A), and we identified several MER41 elements marked by higher chromatin accessibility in EC individuals relative to HCs. This correlated with higher transcript levels of adjacent immunity genes such as GPR35, GSDMD, and TEPCR1 (Figure 4)**^71–73^**. We previously reported that MER41 elements are frequently bound by the transcription factor STAT1, a key transcriptional activator of the interferon response, and have the hallmarks of interferon-inducible enhancers. CRISPR-Cas9 editing was used in cell lines to demonstrate that a subset of MER41 elements indeed function as enhancers driving the interferon-inducibility of several innate immune genes. However, the specific MER41 loci we identified here as differentially active in ECs have not been tested experimentally for enhancer activity; thus, further work is warranted to confirm their regulatory function.

LTR7 and LTR12C are two other HERV families with increased expression in ECs (Figure 5), whose members have been previously implicated in the activation of HIV-1 restriction factors. For example, a recent study found that an LTR7 element controls the transcription of APOBEC3G and APOBEC3H**^33^**, two cytidine deaminases that inhibit HIV-1 replication**^32,83–87^**. Similarly, LTR12C elements have been reported to be activated upon HIV-1 infection and two of these elements act as promoters for GBP2 and GBP5**^31^**, which encode proteins working together to inhibit furin-mediated processing of the HIV-1 envelope and other viral glycoproteins**^30^**. Our work expands upon this seminal discovery by showing that chromatin at these two LTR12C loci is more accessible in some ECs than in HCs, which correlates with increased GBP2 and GBP5 expression in these individuals (Figure 4). We observed an analogous pattern of increased expression and retroelement accessibility in ECs with lncRNA HCP5, an endogenous retroviral gene composed of the 3’ LTR and partial internal region of an ERV3-16 element (Figure 4)**^75^**. HCP5 harbors multiple single nucleotide variants that are associated with HIV-1 control**^88^**^,**89**^, yet the role of HCP5 expression levels in HIV-1 infection had not previously been established. Here, we find increased accessibility of the ERV3 promoter together with increased expression of HCP5 in ECs compared to HCs, suggesting that HCP5 expression may be important in viral control. Thus, our results reinforce the idea that TEs are important regulators of the human antiviral response and uncover several specific elements that appear to be boosting cellular defenses against HIV-1 in ECs. We acknowledge that these conclusions are drawn from correlative patterns and manipulative experiments are needed to infer causality between chromatin changes at these TEs and increased expression of their target genes.

To our knowledge, our study is unique for integrating TEs to the transcriptomic analysis of CD4^+^ T cells of ECs. A major takeaway of our analyses is that there is substantial heterogeneity across the transcriptomes of the 128 EC samples examined, which can be resolved into four distinct clusters each with unique gene and TE expression profiles (Figure 3A & 5A). Importantly, these clusters are neither driven by the origin or processing sites of the samples nor by the genetic ancestry of the individuals (Figure S4), and they cannot be solely explained by variation in CD4^+^ T cell subtype composition (Figure 3B & S4). The transcriptomic heterogeneity we observe is consistent with the known phenotypic plasticity of ECs. For example, it has been reported that the extent of viremic control and T cell population maintenance vary among ECs. Previous studies have also identified multiple genetic determinants of viral control in a subset of ECs, including multiple HLA-B alleles identified through GWAS**^15,16^** and the well-documented *CCR5Δ32* deletion blocking HIV-1 cellular entry**^90^**. Increased cytotoxicity of CD8^+^ T cells and natural killer cells have also been implicated in the EC phenotype**^91–93^**, along with superior HIV-1 antibody and cytokine patterns**^94,95^**. Finally, genomic profiling of EC reservoir T cells has identified unique proviral integration patterns in which defective proviruses are preferentially retained in permissive, euchromatin-rich genomic regions while functional proviruses are relegated to heterochromatin-rich regions**^9,12,18,19^**. Thus, no single factor can universally explain the EC phenotype and one can expect that multiple factors will often combine within an individual to drive HIV-1 resistance. For example, a recent study noted that the *CCR5Δ32* allele occurred in a heterozygous state in only one out of five ECs, while no individual in that cohort held the homozygous deletion**^39^**. Similarly, CCR5 downregulation, another proposed mechanism for elite control of HIV-1**^39^**, was only found to be a defining feature of Cluster 2 in our analysis (Figure 3D). Thus, our data are useful in defining a minimum of four major “transcriptome types’’ that may reflect different combinations of resistance mechanisms, including newly recognized mechanisms involving retroelements. Notably, we found that distinct sets of innate immune genes and restriction factors are upregulated in different clusters, suggesting that elevated basal expression of these factors plays a previously underappreciated role in the elite control phenotype. Further studies will be necessary to cement this idea. We also acknowledge that our study is limited by the small number of ECs individuals with available sequencing data, which likely also limited our ability to identify significant relationships between transcriptome clustering and available patient metadata (Figure S4). While the rarity of ECs in the seropositive population makes it challenging to study this phenotype, the transcriptomic heterogeneity revealed by our analyses underscores the need for surveying larger and more diverse EC cohorts.

Another outstanding question is whether the gene and TE signatures revealed by our analysis of ECs may pre-exist in the general population and are independent of HIV-1 infection or are the result, at least in part, of the initial infection. More broadly, it would be interesting to explore if transcriptomic and/or epigenomic TE profiles can serve as predictive variables for whether an individual will display enhanced viral control. In a recent study investigating the contribution of TEs to variable immune responses to influenza infection, a strong inverse correlation was found between the total amount of pre-infection TE transcripts and post-infection viral load**^96^**. When integrated into a predictive model, the activity of TEs, KZFPs, and KZFP-recruited SETDB1 improved its ability to forecast infection success. These findings suggest their pre-infection TE profiles may be predictive of immune potency. Thus, further studies are needed to explore whether the TE and gene expression profiles we identified in some ECs can be discerned in the general population and used to predict HIV-1 susceptibility. TE profiles may also be predictive of other rare disease trajectories, including the inverse of the EC phenotype: HIV-1 rapid progressors**^97,98^**.

Our study further suggests that the epigenetic state of TEs is a key factor modulating their activity as alternative promoters or enhancers for proximal genes. Prior studies have documented this relationship by comparing the epigenetic states (e.g., DNA methylation, chromatin accessibility) of TE loci across healthy tissues**^28,99,100^** or between cancer and non-cancer cells**^101–104^**, but very few studies have explored this relationship across human individuals. The effects of interindividual epigenetic variation at TEs (metastable epialleles) on adjacent gene expression are well documented in mice and best exemplified by the *viable yellow agouti* allele, which is controlled by a retroviral LTR variably methylated across individuals**^105,106^**. But what controls this type of epigenetic variability? There is growing evidence that KZNF proteins are important trans-regulators of these effects through their ability to bind to TEs in a sequence-specific fashion and recruit the co-repressor KAP1, which in turn recruits chromatin modifying factors like histone and DNA methyltransferases to nucleate local heterochromatin at the bound TEs**^27,78^**. Indeed, recent studies show that genetic variation within TE sequences and/or variation in KZNF expression levels result in variation in chromatin accessibility and *cis*-regulatory activity at different TE loci, with occasional effects on adjacent gene expression**^81,107,108^**.

Motivated by these recent insights, we explored the potential role of KZNFs in mediating the variable expression and chromatin states of TEs among ECs. We found a striking anti-correlation between the expression level of several KZNFs and the TE subfamilies they are known to target (Figure 5C & S6A). For instance, transcript levels of ZNF534 and ZNF430 were strongly anticorrelated with that of HERVH/LTR7 and THE1B elements, respectively (Figure 5C), which are the genomic targets of these KZNFs as determined by ChIP-exo (Figure S6B)**^51,80,81,109,110^**. Notably, the expression of most of these KZNFs is lower in ECs compared to VPs (Figure S6C). These observations suggest that interindividual variation in KZNF expression in CD4^+^ T cells could explain why certain TEs are variably expressed and accessible among ECs. Thus far, there have been few investigations of TE-KZNF interactions in T cells and of their contribution to immunity**^111–113^**. Our results point to a direct interplay between TEs and KZNFs in CD4^+^ T cells, which may have important implications for T cell biology and immune responses. Further work is needed to validate TE-KZNF regulatory interactions in T cells, probe their connection to epigenetic variation at individual TE loci, and explore their repercussions on gene expression variation in CD4^+^ T cells, with and without HIV-1 infection. In sum, our study leads to a testable model (Figure 6) that TEs are regulatory agents contributing to HIV-1 elite control by altering the expression of host immune genes in T cells, and points to KZNFs as important controllers of these activities. As KZNFs are single-copy genes often exclusively dedicated to TE regulation, they represent attractive targets for manipulating TEs for research and therapeutic prospects.

## Supplemental Figures

**Supplementary Figure 1.**
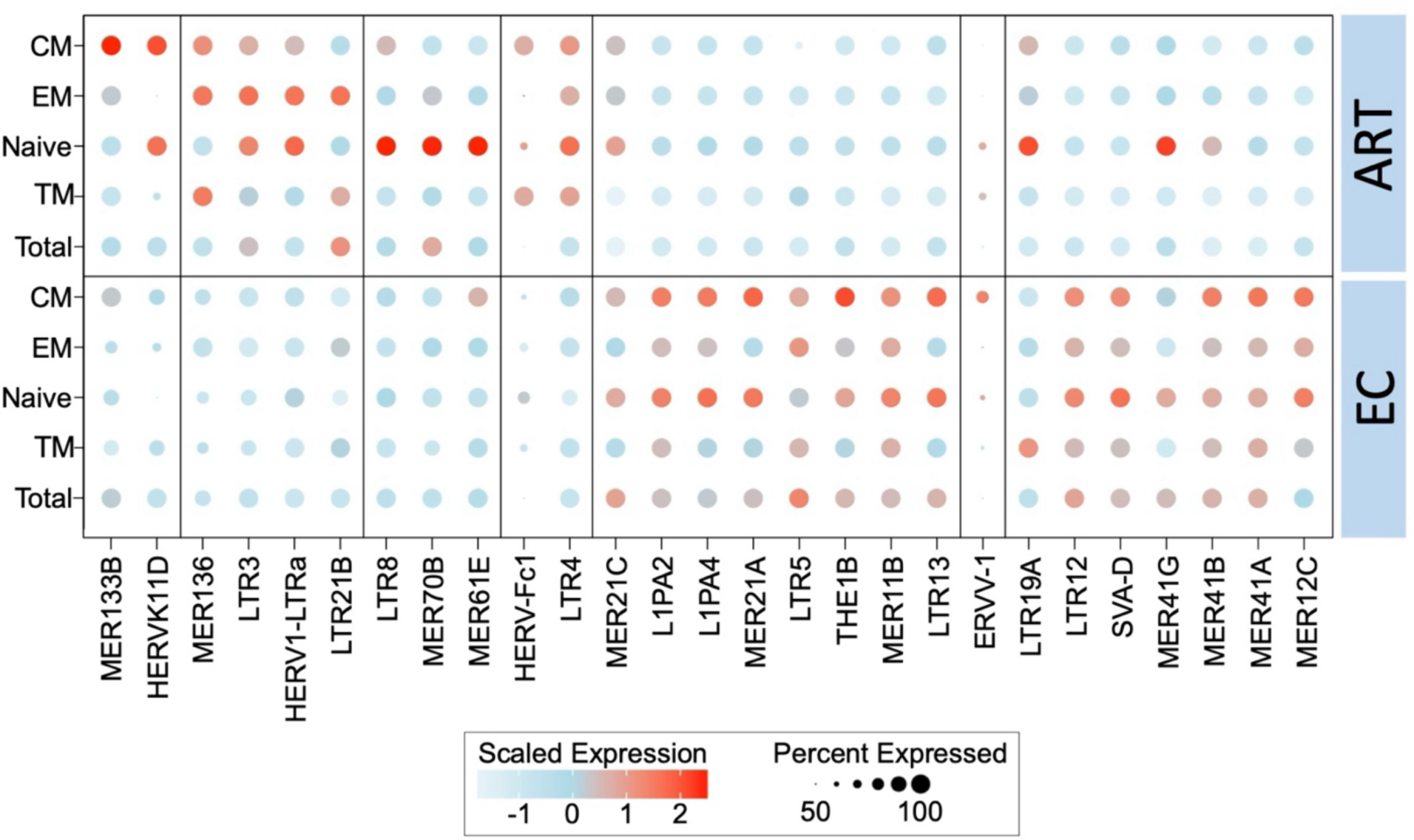
Unique retrotranscriptome of ECs vs PLWH-on-ART. Dot plot illustrating the intensity and abundance of the differential expression of TE families between ECs and PLWH-on-ART, separated by CD4^+^ T cell subtype. Coloration is scaled from lower (white) to higher (red) expression. Point size is directly proportional to the percentage of that sample group expressing the given TE family. Data source: <u>Jiang et al., 2020</u>.

**Supplementary Figure 2.**
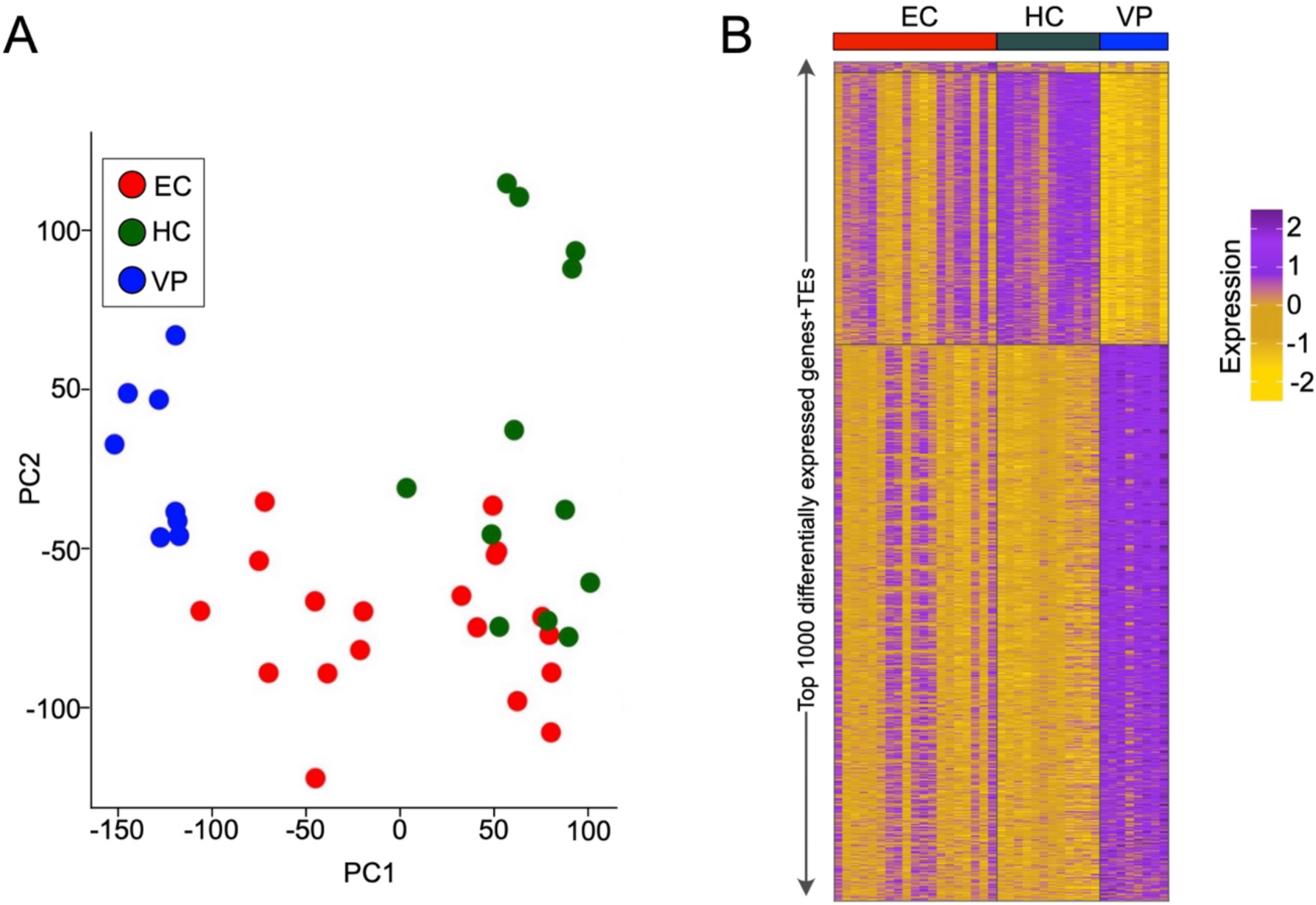
Heterogeneity of EC (retro)transcriptome. B. PCA plot from PBMCs of ECs (red), VPs (blue), and HCs (green), based on the most variably expressed genes and TE families. Every dot is a PBMC sample from an individual. Data source: <u>Zhang et al., 2018</u>. C. Heatmap displaying the scaled expression of genes and TEs distinguishing EC, HC, and VP samples shown in the previous plot. Every row denotes a gene or retroelement. Data source: <u>Zhang et al., 2018</u>.

**Supplementary Figure 3.**
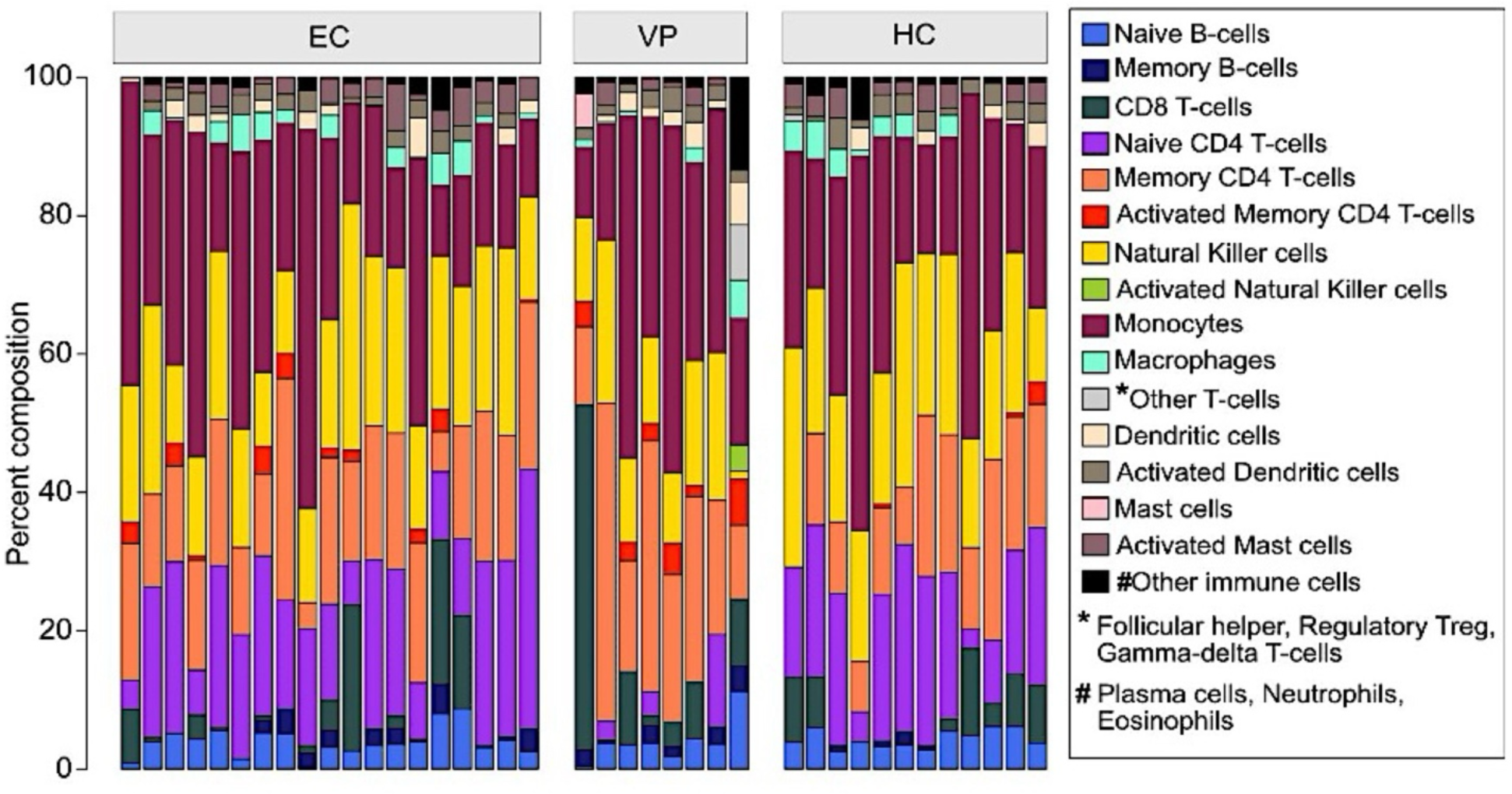
Immune cell profiles of ECs, VPs, and HCs. Stacked barplot showing the immune cell composition of ECs, VPs, and HCs from the deconvolution analysis of PBMC RNA-seq data. Data source: <u>Zhang et al., 2018</u>.

**Supplementary Figure 4.**
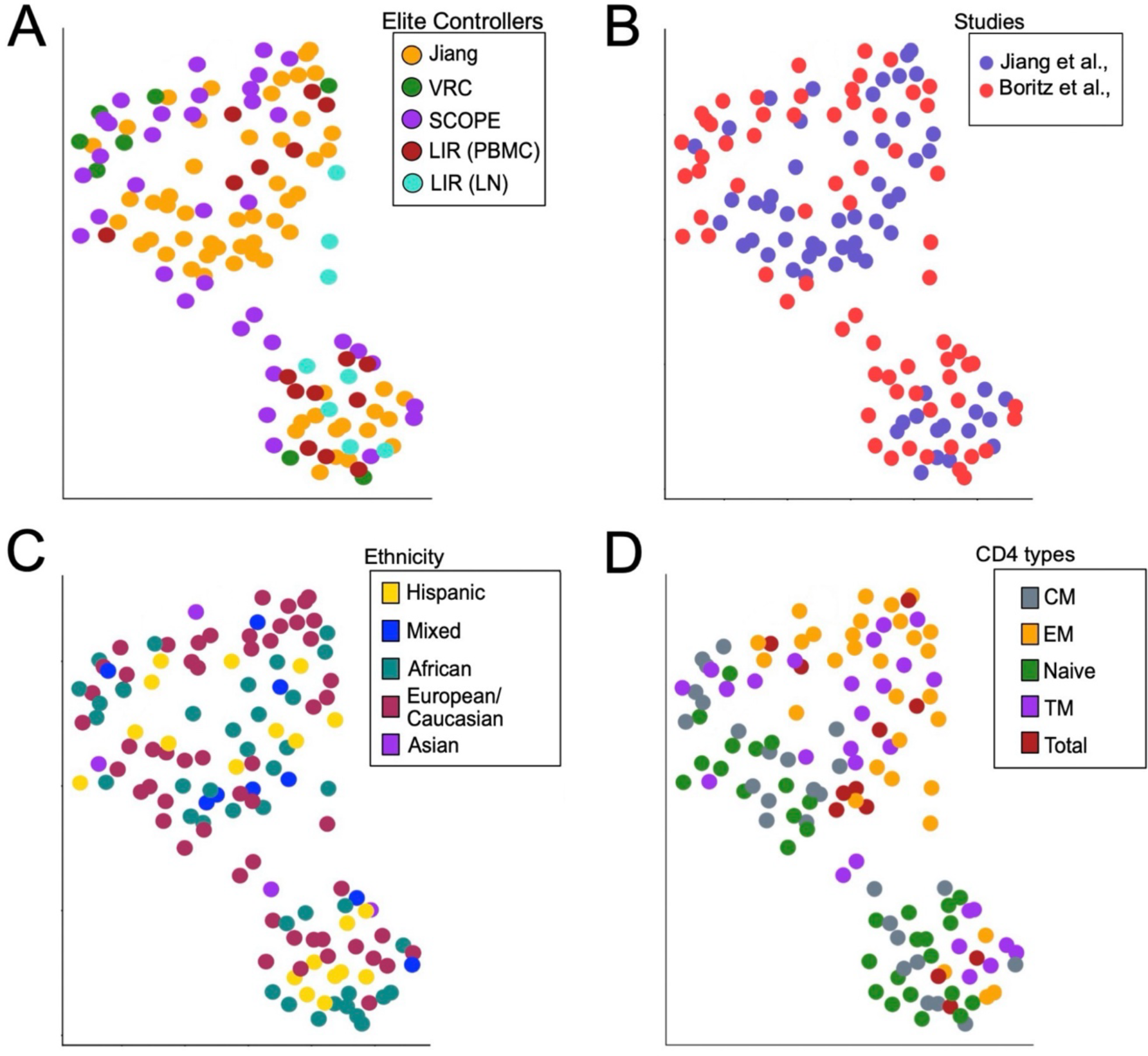
Characterization of the EC clusters. **A.** UMAP plot from Figure 3A with samples color coded based on recruitment location of each patient. Data sources: <u>Jiang et al., 2020</u> and <u>Boritz et al., 2016.</u> **B.** UMAP plot from Figure 3A with samples color coded based on the corresponding source study. Data sources: <u>Jiang et al., 2020</u> and <u>Boritz et al., 2016.</u> **C.** UMAP plot from Figure 3A with samples color coded based on patient ancestry, inferred by HapMap to make direct variant comparisons. Data sources: <u>Jiang et</u> <u>al., 2020</u> and <u>Boritz et al., 2016.</u> **D.** UMAP plot from Figure 3A with samples color coded based on the CD4^+^ subtypes: naïve, central memory (CM), transitional memory (TM), effector memory (EM), and total. Data sources: <u>Jiang et al., 2020</u> and <u>Boritz et al., 2016.</u>

**Supplementary Figure 5.**
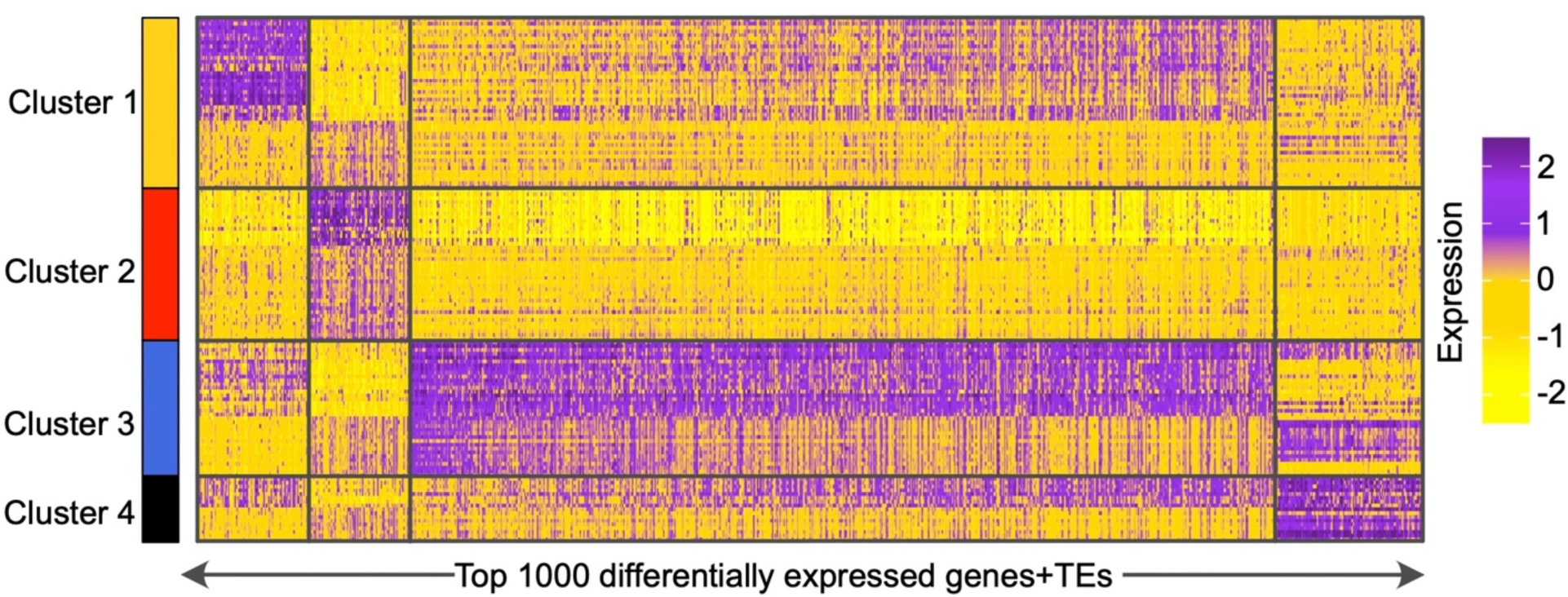
Broad (retro)transcriptomic profile of the EC clusters. Heatmap of the differentially expressed genes (DEGs) that distinguish the EC clusters from each other. Data source: <u>Jiang et al., 2020</u> and <u>Boritz et al., 2016.</u>

**Supplementary Figure 6.**
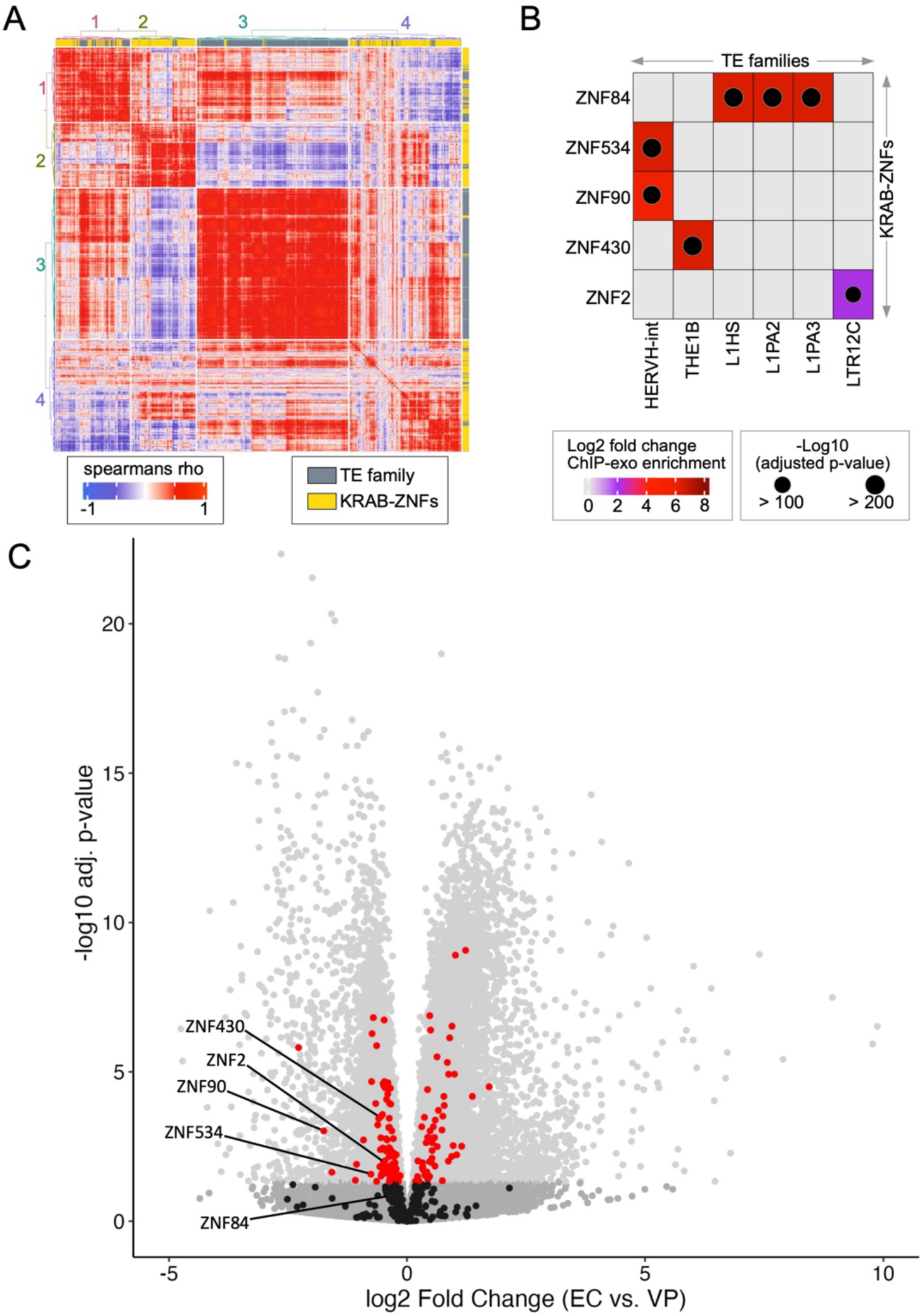
KZNF dynamics in ECs. **A.** Heatmap showing the correlation matrix of TE families with KZNFs across the analyzed EC samples. Clustered pairwise correlation matrix generated by weighted gene co-expression network analysis, using the Spearman’s rank correlation & Euclidean distance. Data sources: <u>Jiang et al., 2020</u> and <u>Boritz et</u> <u>al., 2016.</u> **B.** Heatmap showing the enrichment for binding of a subset of KZNFs to TE families with increased expression in ECs. Coloration is scaled from lower (gray) to higher (red) ChIP-exo enrichment in HEK cells. Point size is directly proportional to the −log10-transformed adjusted p-value. Data source: <u>Imbeault et al., 2017</u>. **C.** Volcano plot displaying the differential expression of genes between EC and VP samples. KZFP genes are in red (p-value < 0.05) or black (p-value > 0.05). All other genes are in gray. Data source: <u>Zhang et al., 2018</u>.

## Acknowledgments

This work was supported in part by a Cornell Presidential Postdoctoral Fellowship to MS and funds from the National Institutes of Health to CF (U01HG009391; R35GM122550), to DFN and CF (UM1AI164559), to DFN (R01CA260691), and a seed grant from the Cornell Office of Academic Integration to CF and DFN. MLB is supported in part by the Department of Medicine Fund for the Future program at Weill Cornell Medicine, sponsored by the Elsa Miller Foundation.

We would like to thank Jez Marston, Melanie Ott, Susana Valente, Lishomwa Ndhlovu, Warner Greene, Ulrike Lange, Zichong Li, Miguel de Mulder Rougvie, Nicholas Dopkins, and members of the Nixon and Feschotte labs for their helpful notes and discussions. We also extend our sincere appreciation to Eli A. Boritz, Daniel C. Douek, Chenyang Jiang, and Xu G. Yu – whose sequencing data was utilized in our analyses – for their open communication throughout this study.

## Author contributions

M.S., S.M.L., L.P.I., M.L.B., D.F.N., and C.F. designed the study. M.S., S.M.L., D.F.N., and C.F. wrote the manuscript. M.S., L.P.I., S.M.L., and M.L.B. performed the data analysis. All authors edited and approved the manuscript.

## Methods

### Data accessibility

All sequencing data is publicly available and can be found in NCBI repositories under the accession numbers GSE122323, PRJNA420459, GSE144334, and GSE83482. The relevant scripts from our analyses are available on <u>our GitHub repository</u>.

### RNA-seq data processing

We mined published RNA-seq datasets of human immune cells isolated from EC, HC, PLWH-on-ART, and VP individuals ranging from PBMCs to CD4^+^ T cell subsets. Quality control was conducted with <u>FastQC</u> on the raw fastq sequencing files. Two nucleotides were removed from the ends based on highly variable quality scores at these locations compared to the rest of the sequence in the RNA-seq reads. Before aligning the resulting reads, we first curated the reference genome annotations using the TE classification from <u>RepeatMasker</u> and the hg19 gene annotation (<u>gencode</u>). We extracted the gene (coding sequences + UTRs) and locus-level TE genomic sequences, combining them to generate a reference transcriptome. These sequences were appended in fasta format. We then annotated every fasta sequence with their respective gene or TE family ID. To guide the transcriptome assembly, we modeled the appended sequences in gtf format, which was then used for expression quantification. Next, we indexed the concatenated gene and TE genomic sequences using ‘salmon’. Finally, we used STAR**^114^** to align the trimmed sequencing reads against our curated hg19 reference genome. The salmon**^115^** tool quantified the counts and normalized expression (transcripts per million; TPM) for each single cell RNA-seq sample. Overall, this approach enabled us to simultaneously calculate TE family and gene expression using expectation-maximization (EM) algorithms. Data integration of the obtained count matrix, normalization at logarithmic scale, and scaling were performed. DESeq2**^116^** was used to perform differential expression analysis between populations.

### Clustering of EC RNA-seq samples

For a subset of EC samples for which differentiated CD4^+^ T cell data was available, we clustered the scaled data by feeding the first five principal components to the graph based KNN method (see GitHub for details). We calculated differential expression and tested their significance level using the Kruskal–Wallis test by comparing the cluster of interest with the rest of the clusters. The Benjamini-Hochberg method further adjusted the obtained p-values to determine the adjusted false discovery rate (FDR). All statistics and visualization of RNA-seq data were performed using <u>R</u>.

### ATAC-seq data processing

The ATAC-seq reads were aligned to the hg19 reference genome by Bowtie 2 (version 2.2.2)**^117^** under the parameters *--very-sensitive-local*. All unmapped reads, non-uniquely mapped reads and PCR duplicates were removed. MACS2**^118^** was used for peak calling. For downstream analysis, we normalized the read counts by computing the numbers of reads per kilobase of bin per million of reads sequenced (RPKM). RPKM values were then averaged for each bin across replicates. To minimize batch and cell type variation, the RPKM values were further normalized by *Z*-score transformation. To visualize the ATAC-seq signal in the UCSC genome browser**^119^**, we extended each read by 100 bp and counted the coverage for each base. To compare ATAC-seq signal over specific TE families between EC clusters, the genome-wide signal was concatenated per cluster and then normalized as mappable reads per million per 100 bp bins. We then visualized each family of interest within a +/- 5 kb genomic window at the elements’ left boundaries according to the hg19 RepeatMasker annotation.

### Multiomic annotation of putative *cis*-regulatory TEs

To identify TEs with putative *cis*-regulatory potential in ECs, we first used intersectBed from bedtools**^120^** with default parameters to calculate the overlap between the ATAC-seq peaks identified in ECs and HCs with the locations of annotated repeats in the hg19 RepeatMasker annotation. The summit output files provided by MACS2 were used to establish a 50% overlapping threshold between each ATAC-seq peak and TE locus. Before the *cis-*regulatory element annotation, we calculated the enrichment of ATAC-seq peaks within each TE family for each population (EC and HC). As repeats of different classes vary greatly in numbers, a random set of ATAC-seq peaks was used for the same analysis as a control. Random peaks were generated by selecting random regions in the genome with the sizes matching each individual TE-associated ATAC-seq peak. The number of observed peaks that overlap with retroelements was compared to the number of random peaks exhibiting the same overlap, and a log2-transformed ratio value was generated as the ‘observed/expected’ enrichment. This TE family-wide significance score was calculated using the R package <u>enrichR</u> while employing the in-house codes (see GitHub for details). Finally, we focused on TE ATAC-seq peaks within 10 kb of annotated transcription start sites (TSS) in hg19. Among these, the peaks which were differentially accessible in ECs compared to HCs (ATAC-seq) and for which the corresponding gene was differentially expressed in ECs compared to HCs (RNA- seq), were annotated as putative EC-specific *cis-*regulatory TE-gene pairs.

## References

1. The path that ends AIDS: UNAIDS Global AIDS Update 2023. (n.d.). Retrieved November 8, 2023, from https://www.unaids.org/en/resources/documents/2023/global-aids-update-2023

2. HIV and AIDS. (n.d.). Retrieved May 31, 2023, from https://www.who.int/news-room/fact-sheets/detail/hiv-aids

3. Global HIV & AIDS statistics—Fact sheet. (n.d.). Retrieved January 19, 2023, from https://www.unaids.org/en/resources/fact-sheet

4. Research priorities for an HIV cure: International AIDS Society Global Scientific Strategy 2021 | Nature Medicine. (n.d.). Retrieved August 18, 2023, from https://www.nature.com/articles/s41591-021-01590-5

5. Hütter, G., Nowak, D., Mossner, M., Ganepola, S., Müssig, A., Allers, K., Schneider, T., Hofmann, J., Kücherer, C., Blau, O., Blau, I. W., Hofmann, W. K., & Thiel, E. (2009). Long-term control of HIV by CCR5 Delta32/Delta32 stem-cell transplantation. The New England Journal of Medicine, 360(7), 692–698. 10.1056/NEJMoa0802905

6. Gupta, R. K., Abdul-jawad, S., McCoy, L. E., Mok, H. P., Peppa, D., Salgado, M., Martinez-Picado, J., Nijhuis, M., Wensing, A. M. J., Lee, H., Grant, P., Nastouli, E., Lambert, J., Pace, M., Salasc, F., Monit, C., Innes, A., Muir, L., Waters, L.,… Olavarria, E. (2019). HIV-1 remission following CCR5Δ32/Δ32 haematopoietic stem cell transplantation. Nature, 568(7751), 244–248. 10.1038/s41586-019-1027-4

7. Psomas, C. K., & Kinloch, S. (n.d.). Highlights of the Conference on Retroviruses and Opportunistic Infections, 4–9 March 2019, Seattle, WA, USA. Journal of Virus Eradication, 5(2), 125–131.

8. Turk, G., Seiger, K., Lian, X., Sun, W., Parsons, E. M., Gao, C., Rassadkina, Y., Polo, M. L., Czernikier, A., Ghiglione, Y., Vellicce, A., Varriale, J., Lai, J., Yuki, Y., Martin, M., Rhodes, A., Lewin, S. R., Walker, B. D., Carrington, M.,… Yu, X. G. (2022). A Possible Sterilizing Cure of HIV-1 Infection Without Stem Cell Transplantation. Annals of Internal Medicine, 175(1), 95–100. 10.7326/L21-0297

9. Jiang, C., Lian, X., Gao, C., Sun, X., Einkauf, K. B., Chevalier, J. M., Chen, S. M. Y., Hua, S., Rhee, B., Chang, K., Blackmer, J. E., Osborn, M., Peluso, M. J., Hoh, R., Somsouk, M., Milush, J., Bertagnolli, L. N., Sweet, S. E., Varriale, J. A.,… Yu, X. G. (2020). Distinct viral reservoirs in individuals with spontaneous control of HIV- Nature, 585(7824), 261–267. 10.1038/s41586-020-2651-8

10. Lifson, A. R., Rutherford, G. W., & Jaffe, H. W. (1988). The natural history of human immunodeficiency virus infection. The Journal of Infectious Diseases, 158(6), 1360–1367. 10.1093/infdis/158.6.1360

11. Navarrete-Muñoz, M. A., Restrepo, C., Benito, J. M., & Rallón, N. (2020). Elite controllers: A heterogeneous group of HIV-infected patients. Virulence, 11(1), 889–897. 10.1080/21505594.2020.1788887

12. Woldemeskel, B. A., Kwaa, A. K., & Blankson, J. N. (2020). Viral reservoirs in elite controllers of HIV-1 infection: Implications for HIV cure strategies. EBioMedicine, 62, 103118. 10.1016/j.ebiom.2020.103118

13. Casado, C., Galvez, C., Pernas, M., Tarancon-Diez, L., Rodriguez, C., Sanchez-Merino, V., Vera, M., Olivares, I., De Pablo-Bernal, R., Merino-Mansilla, A., Del Romero, J., Lorenzo-Redondo, R., Ruiz-Mateos, E., Salgado, M., Martinez-Picado, J., & Lopez-Galindez, C. (2020). Permanent control of HIV-1 pathogenesis in exceptional elite controllers: A model of spontaneous cure. Scientific Reports, 10(1), Article 1. 10.1038/s41598-020-58696-y

14. Gaiha, G. D., Rossin, E. J., Urbach, J., Landeros, C., Collins, D. R., Nwonu, C., Muzhingi, I., Anahtar, M. N., Waring, O. M., Piechocka-Trocha, A., Waring, M., Worrall, D. P., Ghebremichael, M. S., Newman, R. M., Power, K. A., Allen, T. M., Chodosh, J., & Walker, B. D. (2019). Structural topology defines protective CD8+ T cell epitopes in the HIV proteome. Science (New York, N.Y.), 364(6439), 480–484. 10.1126/science.aav5095

15. Rotger, M., Dang, K. K., Fellay, J., Heinzen, E. L., Feng, S., Descombes, P., Shianna, K. V., Ge, D., Günthard, H. F., Goldstein, D. B., Telenti, A., & Immunology, T. S. H. C. S. and the C. for H. V. (2010). Genome-Wide mRNA Expression Correlates of Viral Control in CD4+ T-Cells from HIV-1-Infected Individuals. PLOS Pathogens, 6(2), e1000781. 10.1371/journal.ppat.1000781

16. International HIV Controllers Study, Pereyra, F., Jia, X., McLaren, P. J., Telenti, A., de Bakker, P. I. W., Walker, B. D., Ripke, S., Brumme, C. J., Pulit, S. L., Carrington, M., Kadie, C. M., Carlson, J. M., Heckerman, D., Graham, R. R., Plenge, R. M., Deeks, S. G., Gianniny, L., Crawford, G.,… Zhao, M. (2010). The major genetic determinants of HIV-1 control affect HLA class I peptide presentation. Science (New York, N.Y.), 330(6010), 1551–1557. 10.1126/science.1195271

17. Carrington, M., Bashirova, A. A., & McLaren, P. J. (2013). On stand by: Host genetics of HIV control. AIDS (London, England), 27(18), 2831–2839. 10.1097/01.aids.0000432536.85335.c8

18. Lian, X., Gao, C., Sun, X., Jiang, C., Einkauf, K. B., Seiger, K. W., Chevalier, J. M., Yuki, Y., Martin, M., Hoh, R., Peluso, M. J., Carrington, M., Ruiz-Mateos, E., Deeks, S. G., Rosenberg, E. S., Walker, B. D., Lichterfeld, M., & Yu, X. G. (2021). Signatures of immune selection in intact and defective proviruses distinguish HIV-1 elite controllers. Science Translational Medicine, 13(624), eabl4097. 10.1126/scitranslmed.abl4097

19. Einkauf, K. B., Osborn, M. R., Gao, C., Sun, W., Sun, X., Lian, X., Parsons, E. M., Gladkov, G. T., Seiger, K. W., Blackmer, J. E., Jiang, C., Yukl, S. A., Rosenberg, E. S., Yu, X. G., & Lichterfeld, M. (2022). Parallel analysis of transcription, integration, and sequence of single HIV-1 proviruses. Cell, 185(2), 266–282.e15. 10.1016/j.cell.2021.12.011

20. Huang, A. S., Ramos, V., Oliveira, T. Y., Gaebler, C., Jankovic, M., Nussenzweig, M. C., & Cohn, L. B. (2021). Integration features of intact latent HIV-1 in CD4+ T cell clones contribute to viral persistence. The Journal of Experimental Medicine, 218(12), e20211427. 10.1084/jem.20211427

21. Bourque, G., Burns, K. H., Gehring, M., Gorbunova, V., Seluanov, A., Hammell, M., Imbeault, M., Izsvák, Z., Levin, H. L., Macfarlan, T. S., Mager, D. L., & Feschotte, A. (2018). Ten things you should know about transposable elements. Genome Biology, 19(1), 199. 10.1186/s13059-018-1577-z

22. Rosspopoff, O., & Trono, D. (2023). Take a walk on the KRAB side. Trends in Genetics, 39(11), 844–857. 10.1016/j.tig.2023.08.003

23. Friedli, M., & Trono, D. (2015). The developmental control of transposable elements and the evolution of higher species. Annual Review of Cell and Developmental Biology, 31, 429–451. 10.1146/annurev-cellbio-100814-125514

24. Chuong, E. B., Elde, N. C., & Feschotte, C. (2017). Regulatory activities of transposable elements: From conflicts to benefits. Nature Reviews Genetics, 18(2), Article 2. 10.1038/nrg.2016.139

25. Frank, J. A., Singh, M., Cullen, H. B., Kirou, R. A., Benkaddour-Boumzaouad, M., Cortes, J. L., Garcia Pérez, J., Coyne, C. B., & Feschotte, C. (2022). Evolution and antiviral activity of a human protein of retroviral origin. Science, 378(6618), 422–428. 10.1126/science.abq7871

26. Feschotte, C. (2008). The contribution of transposable elements to the evolution of regulatory networks. Nature Reviews. Genetics, 9(5), 397–405. 10.1038/nrg2337

27. Cosby, R. L., Chang, N.-C., & Feschotte, C. (2019). Host-transposon interactions: Conflict, cooperation, and cooption. Genes & Development, 33(17–18), 1098– 1116. 10.1101/gad.327312.119

28. Fueyo, R., Judd, J., Feschotte, C., & Wysocka, J. (2022). Roles of transposable elements in the regulation of mammalian transcription. Nature Reviews Molecular Cell Biology, 23(7), Article 7. 10.1038/s41580-022-00457-y

29. Chuong, E. B., Elde, N. C., & Feschotte, C. (2016). Regulatory evolution of innate immunity through co-option of endogenous retroviruses. Science (New York, N.Y.), 351(6277), 1083–1087. 10.1126/science.aad5497

30. Braun, E., Hotter, D., Koepke, L., Zech, F., Groß, R., Sparrer, K. M. J., Müller, J. A., Pfaller, C. K., Heusinger, E., Wombacher, R., Sutter, K., Dittmer, U., Winkler, M., Simmons, G., Jakobsen, M. R., Conzelmann, K.-K., Pöhlmann, S., Münch, J., Fackler, O. T.,… Sauter, D. (2019). Guanylate-Binding Proteins 2 and 5 Exert Broad Antiviral Activity by Inhibiting Furin-Mediated Processing of Viral Envelope Proteins. Cell Reports, 27(7), 2092–2104.e10. 10.1016/j.celrep.2019.04.063

31. Srinivasachar Badarinarayan, S., Shcherbakova, I., Langer, S., Koepke, L., Preising, A., Hotter, D., Kirchhoff, F., Sparrer, K. M. J., Schotta, G., & Sauter, D. (2020). HIV-1 infection activates endogenous retroviral promoters regulating antiviral gene expression. Nucleic Acids Research, 48(19), 10890–10908. 10.1093/nar/gkaa832

32. Mangeat, B., Turelli, P., Caron, G., Friedli, M., Perrin, L., & Trono, D. (2003). Broad antiretroviral defence by human APOBEC3G through lethal editing of nascent reverse transcripts. Nature, 424(6944), 99–103. 10.1038/nature01709

33. Singh, M., Kondrashkina, A. M., Widmann, T. J., Cortes, J. L., Bansal, V., Wang, J., Römer, C., Garcia-Canadas, M., Garcia-Perez, J. L., Hurst, L. D., & Izsvák, Z. (2023). A new human embryonic cell type associated with activity of young transposable elements allows definition of the inner cell mass. PLOS Biology, 21(6), e3002162. 10.1371/journal.pbio.3002162

34. Kyriakou, E., & Magiorkinis, G. (2023). Interplay between endogenous and exogenous human retroviruses. Trends in Microbiology, 31(9), 933–946. 10.1016/j.tim.2023.03.008

35. Jones, R. B., Garrison, K. E., Mujib, S., Mihajlovic, V., Aidarus, N., Hunter, D. V., Martin, E., John, V. M., Zhan, W., Faruk, N. F., Gyenes, G., Sheppard, N. C., Priumboom-Brees, I. M., Goodwin, D. A., Chen, L., Rieger, M., Muscat-King, S., Loudon, P. T., Stanley, C.,… Ostrowski, M. A. (2012). HERV-K-specific T cells eliminate diverse HIV-1/2 and SIV primary isolates. The Journal of Clinical Investigation, 122(12), 4473–4489. 10.1172/JCI64560

36. Liu, C.-H., Grandi, N., Palanivelu, L., Tramontano, E., & Lin, L.-T. (2020). Contribution of Human Retroviruses to Disease Development-A Focus on the HIV- and HERV- Cancer Relationships and Treatment Strategies. Viruses, 12(8), 852. 10.3390/v12080852

37. Boritz, E. A., Darko, S., Swaszek, L., Wolf, G., Wells, D., Wu, X., Henry, A. R., Laboune, F., Hu, J., Ambrozak, D., Hughes, M. S., Hoh, R., Casazza, J. P., Vostal, A., Bunis, D., Nganou-Makamdop, K., Lee, J. S., Migueles, S. A., Koup, R. A.,… Douek, D. C. (2016). Multiple Origins of Virus Persistence during Natural Control of HIV Infection. Cell, 166(4), 1004–1015. 10.1016/j.cell.2016.06.039

38. Zhang, W., Ambikan, A. T., Sperk, M., van Domselaar, R., Nowak, P., Noyan, K., Russom, A., Sönnerborg, A., & Neogi, U. (2018). Transcriptomics and Targeted Proteomics Analysis to Gain Insights Into the Immune-control Mechanisms of HIV-1 Infected Elite Controllers. EBioMedicine, 27, 40–50. 10.1016/j.ebiom.2017.11.031

39. Gonzalo-Gil, E., Rapuano, P. B., Ikediobi, U., Leibowitz, R., Mehta, S., Coskun, A. K., Porterfield, J. Z., Lampkin, T. D., Marconi, V. C., Rimland, D., Walker, B. D., Deeks, S., & Sutton, R. E. (2019). Transcriptional down-regulation of ccr5 in a subset of HIV+ controllers and their family members. eLife, 8, e44360. 10.7554/eLife.44360

40. Walker, W. E., Kurscheid, S., Joshi, S., Lopez, C. A., Goh, G., Choi, M., Barakat, L., Francis, J., Fisher, A., Kozal, M., Zapata, H., Shaw, A., Lifton, R., Sutton, R. E., & Fikrig, E. (2015). Increased Levels of Macrophage Inflammatory Proteins Result in Resistance to R5-Tropic HIV-1 in a Subset of Elite Controllers. Journal of Virology, 89(10), 5502–5514. 10.1128/JVI.00118-15

41. Raposo, R. A. S., Abdel-Mohsen, M., Holditch, S. J., Kuebler, P. J., Cheng, R. G., Eriksson, E. M., Liao, W., Pillai, S. K., & Nixon, D. F. (2013). Increased expression of intrinsic antiviral genes in HLA-B*57-positive individuals. Journal of Leukocyte Biology, 94(5), 1051–1059. 10.1189/jlb.0313150

42. McLaren, P. J., & Fellay, J. (2021). HIV-1 and human genetic variation. Nature Reviews Genetics, 22(10), Article 10. 10.1038/s41576-021-00378-0

43. Foster, T. L., Wilson, H., Iyer, S. S., Coss, K., Doores, K., Smith, S., Kellam, P., Finzi, A., Borrow, P., Hahn, B. H., & Neil, S. J. D. (2016). Resistance of Transmitted Founder HIV-1 to IFITM-Mediated Restriction. Cell Host & Microbe, 20(4), 429–442. 10.1016/j.chom.2016.08.006

44. Chaudhuri, A., Yang, B., Gendelman, H. E., Persidsky, Y., & Kanmogne, G. D. (2008). STAT1 signaling modulates HIV-1–induced inflammatory responses and leukocyte transmigration across the blood-brain barrier. Blood, 111(4), 2062–2072. 10.1182/blood-2007-05-091207

45. Chen, J., He, Y., Zhong, H., Hu, F., Li, Y., Zhang, Y., Zhang, X., Lin, W., Li, Q., Xu, F., Chen, S., Zhang, H., Cai, W., & Li, L. (2023). Transcriptome analysis of CD4+ T cells from HIV-infected individuals receiving ART with LLV revealed novel transcription factors regulating HIV-1 promoter activity. Virologica Sinica, 38(3), 398–408. 10.1016/j.virs.2023.03.001

46. D’Antoni, M. L., Kallianpur, K. J., Premeaux, T. A., Corley, M. J., Fujita, T., Laws, E. I., Ogata-Arakaki, D., Chow, D. C., Khadka, V. S., Shikuma, C. M., & Ndhlovu, L. C. (2019). Lower Interferon Regulatory Factor-8 Expression in Peripheral Myeloid Cells Tracks With Adverse Central Nervous System Outcomes in Treated HIV Infection. Frontiers in Immunology, 10, 2789. 10.3389/fimmu.2019.02789

47. Sugawara, S., Reeves, R. K., & Jost, S. (2022). Learning to Be Elite: Lessons From HIV-1 Controllers and Animal Models on Trained Innate Immunity and Virus Suppression. Frontiers in Immunology, 13, 858383. 10.3389/fimmu.2022.858383

48. Morou, A., Brunet-Ratnasingham, E., Dubé, M., Charlebois, R., Mercier, E., Darko, S., Brassard, N., Nganou-Makamdop, K., Arumugam, S., Gendron-Lepage, G., Yang, L., Niessl, J., Baxter, A. E., Billingsley, J. M., Rajakumar, P. A., Lefebvre, F., Johnson, R. P., Tremblay, C., Routy, J.-P.,… Kaufmann, D. E. (2019). Altered differentiation is central to HIV-specific CD4+ T cell dysfunction in progressive disease. Nature Immunology, 20(8), 1059–1070. 10.1038/s41590-019-0418-x

49. Krishnan, S., Wilson, E. M. P., Sheikh, V., Rupert, A., Mendoza, D., Yang, J., Lempicki, R., Migueles, S. A., & Sereti, I. (2014). Evidence for Innate Immune System Activation in HIV Type 1–Infected Elite Controllers. The Journal of Infectious Diseases, 209(6), 931–939. 10.1093/infdis/jit581

50. Díez-Fuertes, F., De La Torre-Tarazona, H. E., Calonge, E., Pernas, M., Alonso-Socas, M. del M., Capa, L., García-Pérez, J., Sakuntabhai, A., & Alcamí, J. (2019). Transcriptome Sequencing of Peripheral Blood Mononuclear Cells from Elite Controller-Long Term Non Progressors. Scientific Reports, 9, 14265. 10.1038/s41598-019-50642-x

51. Pontis, J., Planet, E., Offner, S., Turelli, P., Duc, J., Coudray, A., Theunissen, T. W., Jaenisch, R., & Trono, D. (2019). Hominoid-Specific Transposable Elements and KZFPs Facilitate Human Embryonic Genome Activation and Control Transcription in Naive Human ESCs. Cell Stem Cell, 24(5), 724–735.e5. 10.1016/j.stem.2019.03.012

52. Tang, W. W. C., Dietmann, S., Irie, N., Leitch, H. G., Floros, V. I., Bradshaw, C. R., Hackett, J. A., Chinnery, P. F., & Surani, M. A. (2015). A Unique Gene Regulatory Network Resets the Human Germline Epigenome for Development. Cell, 161(6), 1453–1467. 10.1016/j.cell.2015.04.053

53. Barnada, S. M., Isopi, A., Tejada-Martinez, D., Goubert, C., Patoori, S., Pagliaroli, L., Tracewell, M., & Trizzino, M. (2022). Genomic features underlie the co-option of SVA transposons as cis-regulatory elements in human pluripotent stem cells. PLoS Genetics, 18(6), e1010225. 10.1371/journal.pgen.1010225

54. Patoori, S., Barnada, S. M., Large, C., Murray, J. I., & Trizzino, M. (2022). Young transposable elements rewired gene regulatory networks in human and chimpanzee hippocampal intermediate progenitors. Development, 149(19), dev200413. 10.1242/dev.200413

55. Trizzino, M., Kapusta, A., & Brown, C. D. (2018). Transposable elements generate regulatory novelty in a tissue-specific fashion. BMC Genomics, 19, 468. 10.1186/s12864-018-4850-3

56. Trizzino, M., Park, Y., Holsbach-Beltrame, M., Aracena, K., Mika, K., Caliskan, M., Perry, G. H., Lynch, V. J., & Brown, C. D. (2017). Transposable elements are the primary source of novelty in primate gene regulation. Genome Research, 27(10), 1623–1633. 10.1101/gr.218149.116

57. Wang, T., Zeng, J., Lowe, C. B., Sellers, R. G., Salama, S. R., Yang, M., Burgess, S. M., Brachmann, R. K., & Haussler, D. (2007). Species-specific endogenous retroviruses shape the transcriptional network of the human tumor suppressor protein p53. Proceedings of the National Academy of Sciences of the United States of America, 104(47), 18613–18618. 10.1073/pnas.0703637104

58. Karttunen, K., Patel, D., Xia, J., Fei, L., Palin, K., Aaltonen, L., & Sahu, B. (2023). Transposable elements as tissue-specific enhancers in cancers of endodermal lineage. Nature Communications, 14(1), 5313. 10.1038/s41467-023-41081-4

59. Ivancevic, A., Simpson, D. M., & Chuong, E. B. (2021). Endogenous retroviruses mediate transcriptional rewiring in response to oncogenic signaling in colorectal cancer (p. 2021.10.28.466196). bioRxiv. 10.1101/2021.10.28.466196

60. Gibbs, R. A., Belmont, J. W., Hardenbol, P., Willis, T. D., Yu, F., Yang, H., Ch’ang, L.-Y., Huang, W., Liu, B., Shen, Y., Tam, P. K.-H., Tsui, L.-C., Waye, M. M. Y., Wong, J. T.-F., Zeng, C., Zhang, Q., Chee, M. S., Galver, L. M., Kruglyak, S.,… Methods Group. (2003). The International HapMap Project. Nature, 426(6968), Article 6968. 10.1038/nature02168

61. Kanehisa, M., Furumichi, M., Sato, Y., Kawashima, M., & Ishiguro-Watanabe, M. (2023). KEGG for taxonomy-based analysis of pathways and genomes. Nucleic Acids Research, 51(D1), D587–D592. 10.1093/nar/gkac963

62. The Gene Ontology Consortium, Aleksander, S. A., Balhoff, J., Carbon, S., Cherry, J. M., Drabkin, H. J., Ebert, D., Feuermann, M., Gaudet, P., Harris, N. L., Hill, D. P., Lee, R., Mi, H., Moxon, S., Mungall, C. J., Muruganugan, A., Mushayahama, T., Sternberg, P. W., Thomas, P. D.,… Westerfield, M. (2023). The Gene Ontology knowledgebase in 2023. Genetics, 224(1), iyad031. 10.1093/genetics/iyad031

63. Harris, R. S., Petersen-Mahrt, S. K., & Neuberger, M. S. (2002). RNA editing enzyme APOBEC1 and some of its homologs can act as DNA mutators. Molecular Cell, 10(5), 1247–1253. 10.1016/s1097-2765(02)00742-6

64. Harris, R. S., Bishop, K. N., Sheehy, A. M., Craig, H. M., Petersen-Mahrt, S. K., Watt, I. N., Neuberger, M. S., & Malim, M. H. (2003). DNA deamination mediates innate immunity to retroviral infection. Cell, 113(6), 803–809. 10.1016/s0092-8674(03)00423-9

65. Duan, L., Ozaki, I., Oakes, J. W., Taylor, J. P., Khalili, K., & Pomerantz, R. J. (1994). The tumor suppressor protein p53 strongly alters human immunodeficiency virus type 1 replication. Journal of Virology, 68(7), 4302–4313.

66. Gualberto, A., & Baldwin, A. S. (1995). P53 and Sp1 interact and cooperate in the tumor necrosis factor-induced transcriptional activation of the HIV-1 long terminal repeat. The Journal of Biological Chemistry, 270(34), 19680–19683. 10.1074/jbc.270.34.19680

67. Abdel-Mohsen, M., Raposo, R. A. S., Deng, X., Li, M., Liegler, T., Sinclair, E., Salama, M. S., Ghanem, H. E. A., Hoh, R., Wong, J. K., David, M., Nixon, D. F., Deeks, S. G., & Pillai, S. K. (2013). Expression profile of host restriction factors in HIV-1 elite controllers. Retrovirology, 10(1), 106. 10.1186/1742-4690-10-106

68. Saag, M., & Deeks, S. G. (2010). How Do HIV Elite Controllers Do What They Do? Clinical Infectious Diseases, 51(2), 239–241. 10.1086/653678

69. Zaunders, J., & van Bockel, D. (2013). Innate and Adaptive Immunity in Long-Term Non-Progression in HIV Disease. Frontiers in Immunology, 4. https://www.frontiersin.org/articles/10.3389/fimmu.2013.00095

70. Liao, Y., Wang, J., Jaehnig, E. J., Shi, Z., & Zhang, B. (2019). WebGestalt 2019: Gene set analysis toolkit with revamped UIs and APIs. Nucleic Acids Research, 47(W1), W199–W205. 10.1093/nar/gkz401

71. Boleij, A., Fathi, P., Dalton, W., Park, B., Wu, X., Huso, D., Allen, J., Besharati, S., Anders, R. A., Housseau, F., Mackenzie, A. E., Jenkins, L., Milligan, Graeme., Wu, S., & Sears, C. L. (2021). G-protein coupled receptor 35 (GPR35) regulates the colonic epithelial cell response to enterotoxigenic Bacteroides fragilis. Communications Biology, 4, 585. 10.1038/s42003-021-02014-3

72. Lao, X., Mei, X., Zou, J., Xiao, Q., Ning, Q., Xu, X., Zhang, C., Ji, L., Deng, S., Lu, B., & Chen, M. (2022). Pyroptosis associated with immune reconstruction failure in HIV-1- infected patients receiving antiretroviral therapy: A cross-sectional study. BMC Infectious Diseases, 22(1), 867. 10.1186/s12879-022-07818-0

73. Tang, S.-W., Ducroux, A., Jeang, K.-T., & Neuveut, C. (2012). Impact of cellular autophagy on viruses: Insights from hepatitis B virus and human retroviruses. Journal of Biomedical Science, 19(1), 92. 10.1186/1423-0127-19-92

74. Fellay, J., Shianna, K. V., Ge, D., Colombo, S., Ledergerber, B., Weale, M., Zhang, K., Gumbs, C., Castagna, A., Cossarizza, A., Cozzi-Lepri, A., De Luca, A., Easterbrook, P., Francioli, P., Mallal, S., Martinez-Picado, J., Miro, J. M., Obel, N., Smith, J. P.,… Goldstein, D. B. (2007). A whole-genome association study of major determinants for host control of HIV-1. Science (New York, N.Y.), 317(5840), 944–947. 10.1126/science.1143767

75. Kulski, J. K. (2019). Long Noncoding RNA HCP5, a Hybrid HLA Class I Endogenous Retroviral Gene: Structure, Expression, and Disease Associations. Cells, 8(5), 480. 10.3390/cells8050480

76. Belshaw, R., Pereira, V., Katzourakis, A., Talbot, G., Pačes, J., Burt, A., & Tristem, M. (2004). Long-term reinfection of the human genome by endogenous retroviruses. Proceedings of the National Academy of Sciences of the United States of America, 101(14), 4894–4899. 10.1073/pnas.0307800101

77. Curty, G., Iñiguez, L. P., Nixon, D. F., Soares, M. A., & de Mulder Rougvie, M. (2021). Hallmarks of Retroelement Expression in T-Cells Treated With HDAC Inhibitors. Frontiers in Virology, 1. https://www.frontiersin.org/articles/10.3389/fviro.2021.756635

78. Yang, P., Wang, Y., & Macfarlan, T. S. (2017). The role of KRAB-ZFPs in transposable element repression and mammalian evolution. Trends in Genetics: TIG, 33(11), 871–881. 10.1016/j.tig.2017.08.006

79. Jacobs, F. M. J., Greenberg, D., Nguyen, N., Haeussler, M., Ewing, A. D., Katzman, S., Paten, B., Salama, S. R., & Haussler, D. (2014). An evolutionary arms race between KRAB zinc-finger genes ZNF91/93 and SVA/L1 retrotransposons. Nature, 516(7530), Article 7530. 10.1038/nature13760

80. Tribolet-Hardy, J. de, Thorball, C. W., Forey, R., Planet, E., Duc, J., Coudray, A., Khubieh, B., Offner, S., Pulver, C., Fellay, J., Imbeault, M., Turelli, P., & Trono, D. (2023). Genetic features and genomic targets of human KRAB-zinc finger proteins. Genome Research. 10.1101/gr.277722.123

81. Imbeault, M., Helleboid, P.-Y., & Trono, D. (2017). KRAB zinc-finger proteins contribute to the evolution of gene regulatory networks. Nature, 543(7646), Article 7646. 10.1038/nature21683

82. Bruno, M., Mahgoub, M., & Macfarlan, T. S. (2019). The Arms Race Between KRAB– Zinc Finger Proteins and Endogenous Retroelements and Its Impact on Mammals. Annual Review of Genetics, 53(1), 393–416. 10.1146/annurev-genet-112618-043717

83. Esnault, C., Heidmann, O., Delebecque, F., Dewannieux, M., Ribet, D., Hance, A. J., Heidmann, T., & Schwartz, O. (2005). APOBEC3G cytidine deaminase inhibits retrotransposition of endogenous retroviruses. Nature, 433(7024), 430–433. 10.1038/nature03238

84. Lecossier, D., Bouchonnet, F., Clavel, F., & Hance, A. J. (2003). Hypermutation of HIV-1 DNA in the absence of the Vif protein. Science (New York, N.Y.), 300(5622), 1112. 10.1126/science.1083338

85. Wang, X., Ao, Z., Chen, L., Kobinger, G., Peng, J., & Yao, X. (2012). The Cellular Antiviral Protein APOBEC3G Interacts with HIV-1 Reverse Transcriptase and Inhibits Its Function during Viral Replication. Journal of Virology, 86(7), 3777– 3786. 10.1128/JVI.06594-11

86. Dang, Y., Siew, L. M., Wang, X., Han, Y., Lampen, R., & Zheng, Y.-H. (2008). Human Cytidine Deaminase APOBEC3H Restricts HIV-1 Replication. The Journal of Biological Chemistry, 283(17), 11606–11614. 10.1074/jbc.M707586200

87. Yang, H., Ito, F., Wolfe, A. D., Li, S., Mohammadzadeh, N., Love, R. P., Yan, M., Zirkle, B., Gaba, A., Chelico, L., & Chen, X. S. (2020). Understanding the structural basis of HIV-1 restriction by the full length double-domain APOBEC3G. Nature Communications, 11(1), Article 1. 10.1038/s41467-020-14377-y

88. Yoon, W., Ma, B.-J., Fellay, J., Huang, W., Xia, S.-M., Zhang, R., Shianna, K. V., Liao, H.-X., Haynes, B. F., & Goldstein, D. B. (2010). A polymorphism in the HCP5 gene associated with HLA-B*5701 does not restrict HIV-1 in vitro. AIDS (London, England), 24(1), 155–157. 10.1097/QAD.0b013e32833202f5

89. Thørner, L. W., Erikstrup, C., Harritshøj, L. H., Larsen, M. H., Kronborg, G., Pedersen, C., Larsen, C. S., Pedersen, G., Gerstoft, J., Obel, N., & Ullum, H. (2016). Impact of polymorphisms in the HCP5 and HLA-C, and ZNRD1 genes on HIV viral load. Infection, Genetics and Evolution, 41, 185–190. 10.1016/j.meegid.2016.03.037

90. Liu, R., Paxton, W. A., Choe, S., Ceradini, D., Martin, S. R., Horuk, R., MacDonald, M. E., Stuhlmann, H., Koup, R. A., & Landau, N. R. (1996). Homozygous defect in HIV-1 coreceptor accounts for resistance of some multiply-exposed individuals to HIV-1 infection. Cell, 86(3), 367–377. 10.1016/s0092-8674(00)80110-5

91. Betts, M. R., Nason, M. C., West, S. M., De Rosa, S. C., Migueles, S. A., Abraham, J., Lederman, M. M., Benito, J. M., Goepfert, P. A., Connors, M., Roederer, M., & Koup, R. A. (2006). HIV nonprogressors preferentially maintain highly functional HIV-specific CD8+ T cells. Blood, 107(12), 4781–4789. 10.1182/blood-2005-12-4818

92. Hersperger, A. R., Martin, J. N., Shin, L. Y., Sheth, P. M., Kovacs, C. M., Cosma, G. L., Makedonas, G., Pereyra, F., Walker, B. D., Kaul, R., Deeks, S. G., & Betts, M. R. (2011). Increased HIV-specific CD8+ T-cell cytotoxic potential in HIV elite controllers is associated with T-bet expression. Blood, 117(14), 3799–3808. 10.1182/blood-2010-12-322727

93. Genovese, L., Nebuloni, M., & Alfano, M. (2013). Cell-Mediated Immunity in Elite Controllers Naturally Controlling HIV Viral Load. Frontiers in Immunology, 4, 86. 10.3389/fimmu.2013.00086

94. Nabi, R., Moldoveanu, Z., Wei, Q., Golub, E. T., Durkin, H. G., Greenblatt, R. M., Herold, B. C., Nowicki, M. J., Kassaye, S., Cho, M. W., Pinter, A., Landay, A. L., Mestecky, J., & Kozlowski, P. A. (2017). Differences in serum IgA responses to HIV-1 gp41 in elite controllers compared to viral suppressors on highly active antiretroviral therapy. PLOS ONE, 12(7), e0180245. 10.1371/journal.pone.0180245

95. Jacobs, E. S., Keating, S. M., Abdel-Mohsen, M., Gibb, S. L., Heitman, J. W., Inglis, H. C., Martin, J. N., Zhang, J., Kaidarova, Z., Deng, X., Wu, S., Anastos, K., Crystal, H., Villacres, M. C., Young, M., Greenblatt, R. M., Landay, A. L., Gange, S. J., Deeks, S. G.,… Norris, P. J. (2017). Cytokines Elevated in HIV Elite Controllers Reduce HIV Replication In Vitro and Modulate HIV Restriction Factor Expression. Journal of Virology, 91(6), e02051–16. 10.1128/JVI.02051-16

96. Chen, X., Pacis, A., Aracena, K. A., Gona, S., Kwan, T., Groza, C., Lin, Y. L., Sindeaux, R., Yotova, V., Pramatarova, A., Simon, M.-M., Pastinen, T., Barreiro, L. B., & Bourque, G. (2023). Transposable elements are associated with the variable response to influenza infection. Cell Genomics, 3(5), 100292. 10.1016/j.xgen.2023.100292

97. McLaren, P. J., & Carrington, M. (2015). The impact of host genetic variation on infection with HIV-1. Nature Immunology, 16(6), 577–583. 10.1038/ni.3147

98. Zhang, Z.-N., Xu, J.-J., Fu, Y.-J., Liu, J., Jiang, Y.-J., Cui, H.-L., Zhao, B., Sun, H., He, Y.-W., Li, Q.-J., & Shang, H. (2013). Transcriptomic analysis of peripheral blood mononuclear cells in rapid progressors in early HIV infection identifies a signature closely correlated with disease progression. Clinical Chemistry, 59(8), 1175–1186. 10.1373/clinchem.2012.197335

99. Xie, M., Hong, C., Zhang, B., Lowdon, R. F., Xing, X., Li, D., Zhou, X., Lee, H. J., Maire, C. L., Ligon, K. L., Gascard, P., Sigaroudinia, M., Tlsty, T. D., Kadlecek, T., Weiss, A., O’Geen, H., Farnham, P. J., Madden, P. A. F., Mungall, A. J.,… Wang, T. (2013). DNA hypomethylation within specific transposable element families associates with tissue-specific enhancer landscape. Nature Genetics, 45(7), 836– 841. 10.1038/ng.2649

100. Ecco, G., Cassano, M., Kauzlaric, A., Duc, J., Coluccio, A., Offner, S., Imbeault, M., Rowe, H. M., Turelli, P., & Trono, D. (2016). Transposable elements and their KRAB-ZFP controllers regulate gene expression in adult tissues. Developmental Cell, 36(6), 611–623. 10.1016/j.devcel.2016.02.024

101. Leung, A., Trac, C., Kato, H., Costello, K. R., Chen, Z., Natarajan, R., & Schones, D. E. (2018). LTRs activated by Epstein-Barr virus–induced transformation of B cells alter the transcriptome. Genome Research, 28(12), 1791–1798. 10.1101/gr.233585.117

102. Babaian, A., Romanish, M. T., Gagnier, L., Kuo, L. Y., Karimi, M. M., Steidl, C., & Mager, D. L. (2016). Onco-exaptation of an endogenous retroviral LTR drives IRF5 expression in Hodgkin lymphoma. Oncogene, 35(19), 2542–2546. 10.1038/onc.2015.308

103. Lock, F. E., Rebollo, R., Miceli-Royer, K., Gagnier, L., Kuah, S., Babaian, A., Sistiaga-Poveda, M., Lai, C. B., Nemirovsky, O., Serrano, I., Steidl, C., Karimi, M. M., & Mager, D. L. (2014). Distinct isoform of FABP7 revealed by screening for retroelement-activated genes in diffuse large B-cell lymphoma. Proceedings of the National Academy of Sciences of the United States of America, 111(34), E3534– E3543. 10.1073/pnas.1405507111

104. Lamprecht, B., Walter, K., Kreher, S., Kumar, R., Hummel, M., Lenze, D., Köchert, K., Bouhlel, M. A., Richter, J., Soler, E., Stadhouders, R., Jöhrens, K., Wurster, K. D., Callen, D. F., Harte, M. F., Giefing, M., Barlow, R., Stein, H., Anagnostopoulos, I.,… Mathas, S. (2010). Derepression of an endogenous long terminal repeat activates the CSF1R proto-oncogene in human lymphoma. Nature Medicine, 16(5), 571–579, 1p following 579. 10.1038/nm.2129

105. Bertozzi, T. M., & Ferguson-Smith, A. C. (2020). Metastable epialleles and their contribution to epigenetic inheritance in mammals. Seminars in Cell & Developmental Biology, 97, 93–105. 10.1016/j.semcdb.2019.08.002

106. Bertozzi, T. M., Becker, J. L., Blake, G. E. T., Bansal, A., Nguyen, D. K., Fernandez-Twinn, D. S., Ozanne, S. E., Bartolomei, M. S., Simmons, R. A., Watson, E. D., & Ferguson-Smith, A. C. (2021). Variably methylated retrotransposons are refractory to a range of environmental perturbations. Nature Genetics, 53(8), 1233–1242. 10.1038/s41588-021-00898-9

107. Coluccio, A., Ecco, G., Duc, J., Offner, S., Turelli, P., & Trono, D. (2018). Individual retrotransposon integrants are differentially controlled by KZFP/KAP1-dependent histone methylation, DNA methylation and TET-mediated hydroxymethylation in naïve embryonic stem cells. Epigenetics & Chromatin, 11(1), 7. 10.1186/s13072-018-0177-1

108. Bertozzi, T. M., Elmer, J. L., Macfarlan, T. S., & Ferguson-Smith, A. C. (2020). KRAB zinc finger protein diversification drives mammalian interindividual methylation variability. Proceedings of the National Academy of Sciences, 117(49), 31290– 31300. 10.1073/pnas.2017053117

109. Carter, T. A., Singh, M., Dumbović, G., Chobirko, J. D., Rinn, J. L., & Feschotte, C. (2022). Mosaic cis-regulatory evolution drives transcriptional partitioning of HERVH endogenous retrovirus in the human embryo. eLife, 11, e76257. 10.7554/eLife.76257

110. Altemose, N., Noor, N., Bitoun, E., Tumian, A., Imbeault, M., Chapman, J. R., Aricescu, A. R., & Myers, S. R. (2017). A map of human PRDM9 binding provides evidence for novel behaviors of PRDM9 and other zinc-finger proteins in meiosis. eLife, 6, e28383. 10.7554/eLife.28383

111. Tie, C. H., Fernandes, L., Conde, L., Robbez-Masson, L., Sumner, R. P., Peacock, T., Rodriguez-Plata, M. T., Mickute, G., Gifford, R., Towers, G. J., Herrero, J., & Rowe, H. M. (2018). KAP1 regulates endogenous retroviruses in adult human cells and contributes to innate immune control. EMBO Reports, 19(10). 10.15252/embr.201745000

112. Turelli, P., Castro-Diaz, N., Marzetta, F., Kapopoulou, A., Raclot, C., Duc, J., Tieng, V., Quenneville, S., & Trono, D. (2014). Interplay of TRIM28 and DNA methylation in controlling human endogenous retroelements. Genome Research, 24(8), 1260– 1270. 10.1101/gr.172833.114

113. Marzetta, F., Simó-Riudalbas, L., Duc, J., Planet, E., Verp, S., Turelli, P., & Trono, D. (2019). The KZFP/KAP1 system controls transposable elements-embedded regulatory sequences in adult T cells (p. 523597). bioRxiv. 10.1101/523597

114. Dobin, A., Davis, C. A., Schlesinger, F., Drenkow, J., Zaleski, C., Jha, S., Batut, P., Chaisson, M., & Gingeras, T. R. (2013). STAR: Ultrafast universal RNA-seq aligner. Bioinformatics (Oxford, England), 29(1), 15–21. 10.1093/bioinformatics/bts635

115. Salmon provides fast and bias-aware quantification of transcript expression | Nature Methods. (n.d.). Retrieved December 11, 2023, from https://www.nature.com/articles/nmeth.4197

116. Love, M. I., Huber, W., & Anders, S. (2014). Moderated estimation of fold change and dispersion for RNA-seq data with DESeq2. Genome Biology, 15(12), 550. 10.1186/s13059-014-0550-8

117. Langmead, B., & Salzberg, S. L. (2012). Fast gapped-read alignment with Bowtie 2. Nature Methods, 9(4), 357–359. 10.1038/nmeth.1923

118. Zhang, Y., Liu, T., Meyer, C. A., Eeckhoute, J., Johnson, D. S., Bernstein, B. E., Nusbaum, C., Myers, R. M., Brown, M., Li, W., & Liu, X. S. (2008). Model-based Analysis of ChIP-Seq (MACS). Genome Biology, 9(9), R137. 10.1186/gb-2008-9-9-r137

119. The Human Genome Browser at UCSC. (n.d.). Retrieved December 11, 2023, from https://genome.cshlp.org/content/12/6/996.abstract

120. Quinlan, A. R., & Hall, I. M. (2010). BEDTools: A flexible suite of utilities for comparing genomic features. Bioinformatics, 26(6), 841–842. 10.1093/bioinformatics/btq033

